# TCR/ITK signaling *via* mTOR tunes CD8^+^ T cell homeostatic proliferation, metabolism, and anti-tumor effector function

**DOI:** 10.1101/359604

**Authors:** Weishan Huang, Junjie Luo, Avery August

## Abstract

T cell homeostatic proliferation (HP) is regulated by T cell receptor (TCR) signals and homeostatic cytokines, and suggested to be proportional to TCR signal strength. However, we show here that ITK, a positive regulator of TCR signaling, negatively tunes CD8^+^ T cell HP, metabolism, and effector function. Under lymphopenic environments, *Itk*^−/−^ CD8^+^ T cells exhibit significant increase in T cell-intrinsic HP, which requires mTOR activity and can be driven by T cell-T cell interaction. TCR signals through ITK tune IL-7-mediated CD8^+^ T cell metabolism and HP in a mTOR-dependent manner. The lack of ITK also resulted in enhanced effector cell fate programming, antigen sensitivity and anti-tumor immunity by HP cells. Thus, TCR signaling via ITK, is a negative tuner of CD8^+^ T cell homeostasis, metabolism and effector function, and may be a target for clinical benefit in cancer therapy.

**One Sentence Summary:** TCR signal strength had been long-thought to be proportional to T cell proliferation and effector function, here we demonstrate a counterintuitive role of the TCR signaling through ITK in negatively tuning proliferation under lymphopenic conditions via regulating mTOR activity, T cell metabolism, proliferation, and effector function.

## Introduction

CD8^+^ T cells are involved in a broad spectrum of immune activities including viral, bacterial and parasitic infections, immune disorders and anti-tumor responses. Antigenic stimulation normally drives the development of short-lived effectors and induces long-lived memory CD8^+^ T cells that can provide effector immunity against the current infection and in some cases, life-long protection against reinfection respectively (1, 2). However, in the unimmunized homeostatic state, a number of other memory-like CD8^+^ T cell types have been described, including virtual memory cells found in normal homeostatic environment, innate memory phenotype cells that can be driven by the cytokine IL-4, and those derived through lymphopenia-induced homeostatic proliferation (HP CD8^+^ T cells) (3–5).

HP CD8^+^ T cells are typically derived from the proliferation of naïve T cells in response to lymphopenic conditions, dependent on cytokines (IL-7 and/or IL-15) and tonic signals from self-antigen-MHCI interactions (6–9). During lymphopenia-induced HP, CD8^+^ T cells undergo expansion and functional conversion, progressively acquiring a memory-like phenotype (5). However, they have distinct features from true memory cells generated by antigenic stimulation. HP CD8^+^ T cells do not up-regulate antigen-inducible T cell early activation markers, or shed CD62L like antigen-experienced effector memory cells (9–11). Furthermore, like antigen-induced true memory cells, HP CD8^+^ T cells require the presence of CD4^+^ T cells during proliferation to acquire protective memory function (12). These CD4-helped HP CD8^+^ T cells have been shown to provide considerable protection against infection (12) and tumor growth (13, 14). Following initiation of this HP, IL-7 is essential in inducing mTOR activity that further induces T-bet expression, which contributes to the formation of protective CD8^+^ T cell function in the latter stages. This later phase of HP-induced memory differentiation requires IL-15, reduced mTOR activity and T-bet expression, and enhanced expression of the related transcription factor Eomesodermin (Eomes) (15). The tonic signals from self-antigen-MHCI interactions that drive HP have been suggested to be positively correlated with the affinity of TCR for self-antigen-MHCI, such that TCRs with low affinity exhibit reduced HP compared to those with higher affinity (16). This has led to the suggestion that the strength of the TCR’s signals is a positive regulator of HP (17, 18).

The Tec family kinase ITK is an essential signaling mediator downstream of T cell receptor (TCR), regulating the strength the TCR signaling (19). ITK plays critical roles in T cell activation and differentiation (for review see (20)). In the mice lacking ITK, CD8^+^ T cells spontaneously develop a memory-like phenotype independent of prior specific antigenic stimulation (21, 22), with high levels of Eomes and the capacity to rapidly produce IFN-γ upon stimulation (21–24). These innate memory-like CD8^+^ T cells in *Itk*^−/−^ mice depend on the IL-4/STAT6 signaling axis for development (3, 4), and are transcriptomically distinct from HP memory-like CD8^+^ T cells (23).

Given the role ITK plays in regulating TCR signal strength, we tested the assumption that HP of CD8^+^ T cells is proportional to the strength of TCR interaction with self antigen-MHCI. We show here that ITK is a key regulator in CD8^+^ T cell HP and the accompanying effector programing. However, contrary to current models, ITK plays a negative role in this process, since *Itk*^−/−^ naïve CD8^+^ T cells exhibited massive and immediate expansion and transition into effector phenotype cells when encountering lymphopenic conditions. TCR activation dampens CD8^+^ T cell HP and IL-7-mediated metabolic activities, which requires the function of ITK. In the absence of ITK, mTORC1 activity contributes to the enhanced expansion, effector cell programming, and IL-7-induced metabolic activity during CD8^+^ T cell homeostasis. Moreover, ITK is involved in retuning the antigen sensitivity that occurs during homeostatic expansion, and suppresses HP effector cell-mediated anti-tumor immunity. Our data suggest that ITK, and TCR signal strength, negatively tunes critical aspects of T cell HP and effector cell development, and that this can be used to target CD8^+^ T cells for enhanced tumor immunotherapy.

## Results

### ITK regulates CD8^+^ T cell homeostatic proliferation under lymphopenic conditions

In order to investigate the role of ITK in naïve CD8^+^ T cell expansion in a lymphopenic environment, we generated *Itk*^−/−^ mice carrying an OTI transgene (transgenic TCR expressed only by CD8^+^ T cells, specific for chicken ovalbumin 257-264 peptide presented by MHCI H-2k^b^ (25)) on the *Rag*^−/−^ background. CD8^+^ T cells in *Itk*^−/−^ OTI-*Rag*^−/−^ mice are predominantly naïve, unlike what is observed in the *Itk*^−/−^ mice (Figure 1A). We purified naïve (CD44^lo^CD62L^hi^CD122^−^) WT and *Itk*^−/−^ CD8^+^ T cells from these mice, and transferred them into lymphopenic recipients including *Rag*^−/−^ (H-2k^b^), *Rag^−/−^γc^−/−^* (H-2k^b^), NSG (different MHC haplotype H-2k^d^), as well as sub-lethally γ-irradiated WT (H-2k^b^) mice, to examine their capacity for homeostatic expansion (Figure 1B & C). Contrary to expectation, we found that significantly larger populations of *Itk*^−/−^ CD8^+^ T cells 10 days post transfer compared to WT cells, in all lymphopenic murine models regardless of MHC haplotype (Figure 1B).

**Figure 1.**
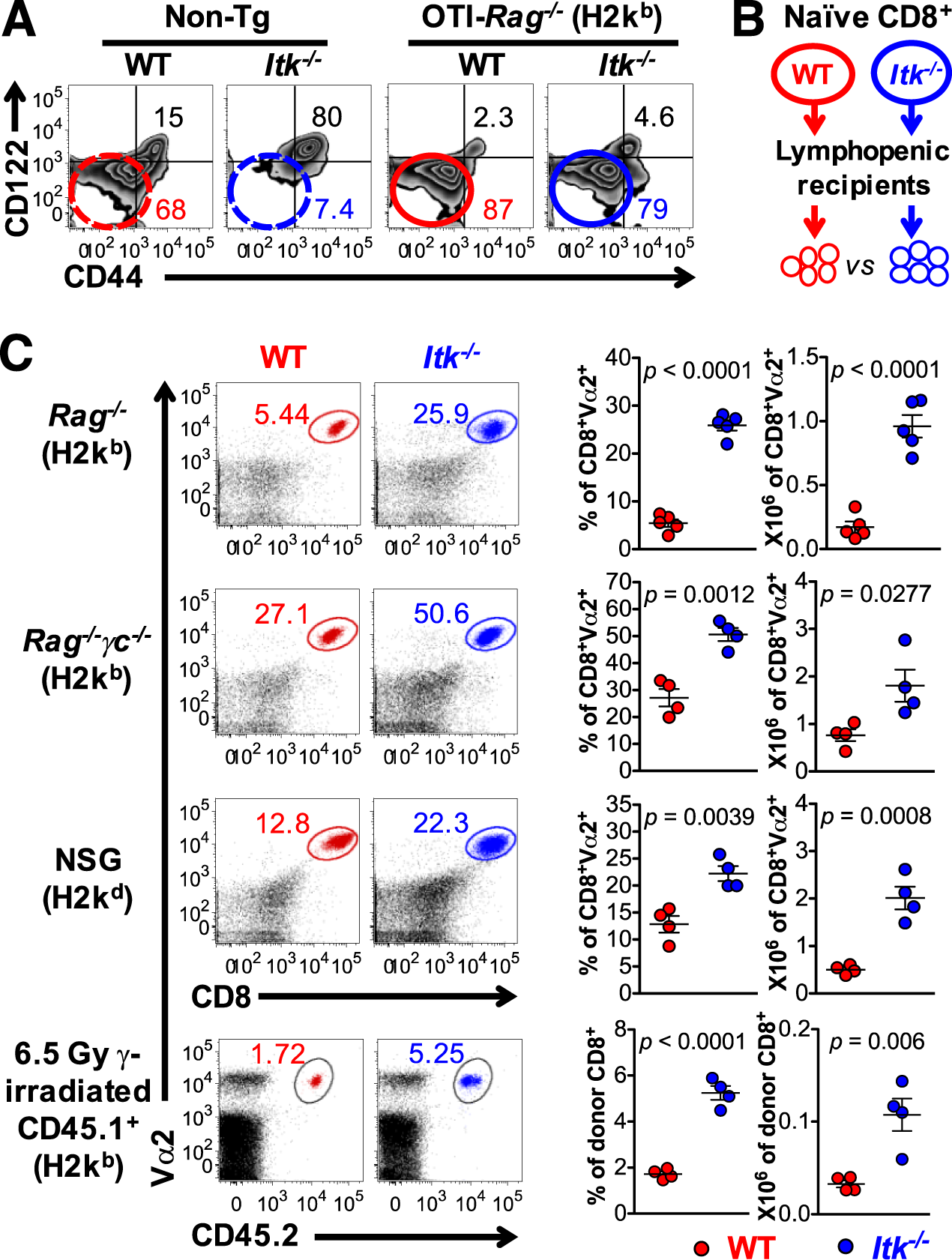
ITK deficiency in naïve CD8^+^ T cells results in massive early expansion during lymphopenia-induced homeostatic expansion. (A) Representative plots of expression of CD44/CD122 on splenic CD8^+^ T cell from non-transgenic (Non-Tg) or OTI-*Rag*^−/−^ transgenic WT and *Itk*^−/−^ mice (haplotype H2K^b^). (B) Experimental model of WT and *Itk*^−/−^ CD8^+^ T cell expansion under lymphopenic environments. Naïve CD8^+^ T cells were flow-sorted from circled area in (A) and 0.5 χ 10^6^ of WT or *Itk*^−/−^ naïve cells (CD45.2^+^) were transferred into lymphopenic *Rag^−/−^, Rag^−/−^yc^−/−^*, NSG, or sublethally irradiated CD45.2-congenic hosts. (C) Representative plots of WT and *Itk*^−/−^ CD8^+^ T cells following 10 days of HP; summary of percentage and number of HP CD8^+^ T cells of viable lymphocytes in the indicated lymphopenic hosts. Total number of donor CD8^+^ T cells in the spleen and lymph node of lymphopenic recipients is shown. Data presented as Mean ± SEM; *p* values generated by non-parametric Mann-Whitney test. N = 4. Data represent results of 3 independent experiments.

### ITK regulation of T cell homeostatic proliferation is T cell-intrinsic

Although WT and *Itk*^−/−^ naïve CD8^+^ T cells started out with the same initial naïve phenotype, and were transferred and initiated lymphopenia-induced HP in the same initial environmental setting, *Itk*^−/−^ cells turned on a massive expansion program resulting in a significantly larger population compared to WT cells (Figure 1). The large *Itk*^−/−^ HP CD8^+^ T cell population at the early phase of lymphopenia-induced proliferation may have a significant influence on the conditions *in vivo*, for example, on the availability of homeostatic cytokines and cell-cell interaction. This may further lead to the altered phenotype observed in *Itk*^−/−^ HP CD8^+^ T cells, including proliferation, death and effector differentiation. To test whether ITK has T cell-intrinsic function in tuning lymphopenia-induced HP, we performed an equally mixed donor transfer utilizing naïve T cells from congenic CD45.1^+^ WT and CD45.2^+^ *Itk*^−/−^ mice on the OTI-*Rag*^−/−^ background as donors (Figure 2A), which allows WT and *Itk*^−/−^ CD8^+^ T cells to proliferate in exactly the same environment along the time course. Again, we found that *Itk*^−/−^ CD8^+^ T cells exhibited the hyperactive expansion response (Figure 2B-C) compared to co-expanding WT cells.

**Figure 2.**
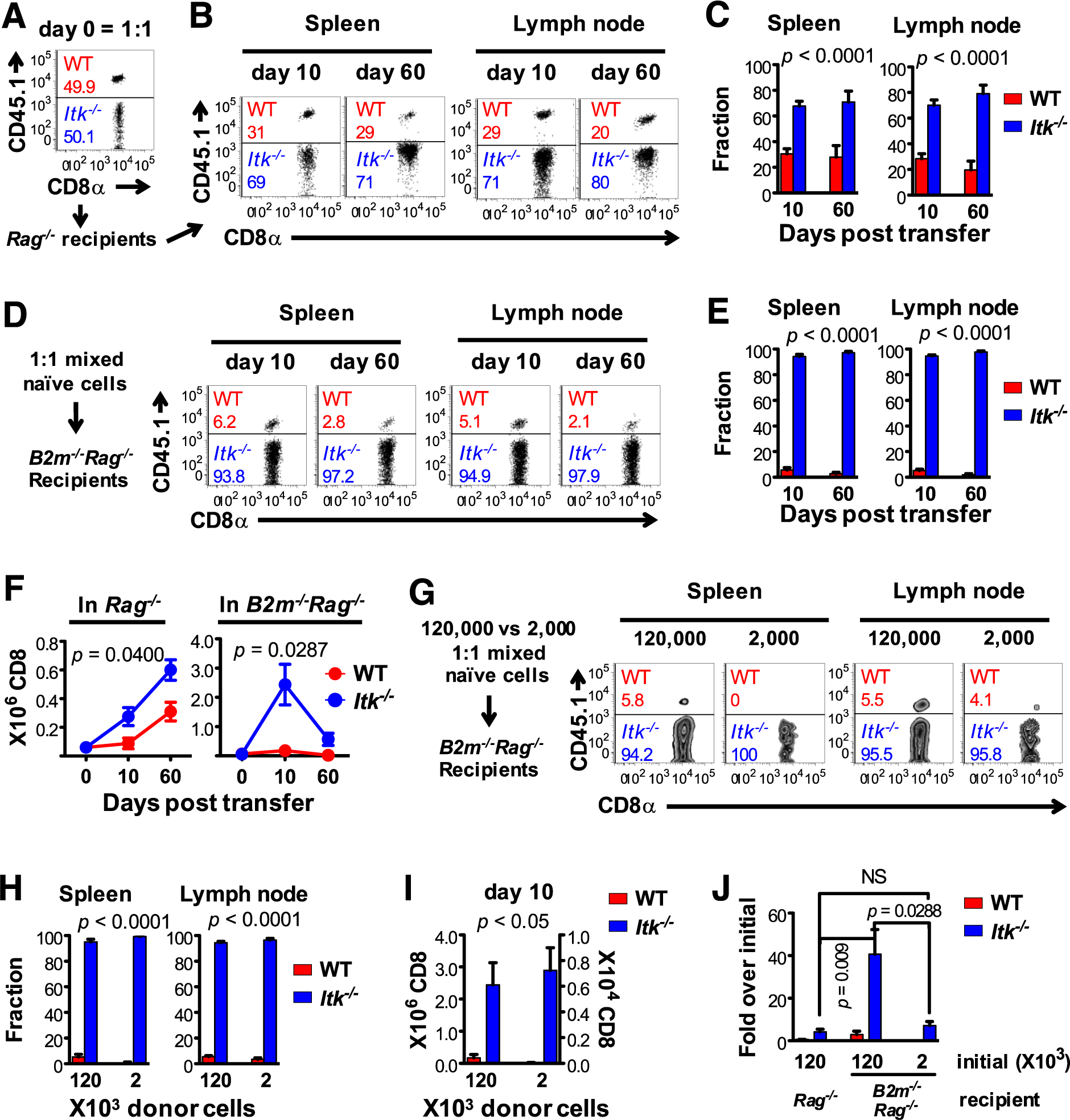
ITK intrinsically regulates CD8^+^ T cell proliferation under lymphopenic conditions independently of B2m/MHCI expression. (A) Purified congenic naïve WT (CD45.1^+^) and *Itk*^−/−^ (CD45.1) OTI-*Rag*^−/−^ CD8^+^ T cells were mixed at the ratio of 1:1, and a total 120,000 T cells were transferred to *Rag*^−/−^ lymphopenic recipients, and analyzed 10 and 60 days post transfer in (B-C). N = 4 - 6, from 2 independent experiments. (B) Representative plots and (C) summary of fractions of WT and *Itk*^−/−^ CD8^+^ T cells expanded in lymphopenic environment. Data represent Mean ± SEM. *p* values generated by non-parametric Mann-Whitney test, comparing WT and *Itk*^−/−^. (D-F) 1:1 mixture of WT and *Itk*^−/−^ naïve OTI-*Rag*^−/−^ CD8^+^ T cells (total 120,000) were transferred into *B2m^−/−^ Rag^−/−^* recipients, and analyzed 10 and 60 days post transfer. N = 3 - 6, from 2 independent experiments. (D) Representative plots and (E) summary of fractions of WT and *Itk*^−/−^ T cells expanded. *p* values generated by non-parametric Mann-Whitney test, comparing WT and *Itk*^−/−^ at each time point. (F) Number of expanded WT and *Itk*^−/−^ CD8^+^ T cells. *p* value generated by two-way ANOVA. (G-I) 1:1 mixed WT and *Itk*^−/−^ naïve *OTI-Rag*^−/−^ CD8^+^ T cells (total 120,000 or 2,000) were transferred into *B2m^−/−^ Rag^−/−^* recipients, and analyzed 10 days post transfer. N ≥ 4, combined from independent experiments. *p* values generated by non-parametric Mann-Whitney test. N = 4 - 5, combined from 2 independent experiments. Data represent Mean ± SEM. (G) Representative plots and (H) summary of fractions of WT and *Itk*^−/−^ T cells expanded in lymphopenic environment. (I) Number of WT and *Itk*^−/−^ CD8^+^ T cells expanded from 120,000 (left y-axis) or 2,000 (right y-axis) initial cells. (J) Fold expansion of 1:1 mixed WT and *Itk*^−/−^ CD8^+^ T cells in the same environment: 120,000 initial cells in *Rag*^−/−^ or *B2m^−/−^Rag^−/−^* recipients, or 2,000 initial cells in *B2m^−/−^Rag^−/−^* recipients. NS = “No Significance”.

### Hyperactive proliferation of Itk^−/−^ T cells independent of recipient MHCI

Lymphopenia-induced CD8^+^ T cell HP has been shown to be independent of *cognate* antigenic simulation, but dependent on tonic interactions between self-antigen-MHC and the TCR (6, 7). However, maintenance of CD8^+^ memory T cells is independent of host MHC interaction with TCR (11). Thus, we considered whether MHCI expression by the lymphopenic recipients is required to tonically stimulate the TCR (and thus ITK) to regulate the hyper-proliferation of *Itk*^−/−^ CD8^+^ T cells. To investigate this, we used *Rag*^−/−^ mice that lack expression of β2 microglobulin (B2m) resulting in lack of cell surface expression of MHCI (*B2m*^−/−^*Rag*^−/−^), as lymphopenic recipients. Analysis of WT or *Itk*^−/−^ CD8^+^ T cells transferred singly into *B2m*^−/−^*Rag*^−/−^ hosts revealed, to our surprise, that recipient MHCI is dispensable for hyper-proliferation of *Itk*^−/−^ CD8^+^ T cells in this lymphopenic environment, as transfer of *Itk*^−/−^ CD8^+^ T cells gave rise to a significantly larger HP CD8+ T cell population in MHCI deficient hosts than WT cells (Figure S1A&B). Furthermore, co-transfer of equal numbers of WT and *Itk*^−/−^ naïve CD8^+^ T cells (120,000 in total) into the same *B2m*^−/−^ *Rag*^−/−^ recipients gave rise to predominantly *Itk*^−/−^ HP CD8^+^ T cells (Figure 2D&E), resulting in an even larger *Itk*^−/−^ HP population compared to transfer into MHCI expressing recipients (Figure 2F). These data suggest that significant tonic interaction between recipient-MHCI and HP CD8+ T cell receptor is not required for *Itk*^−/−^ HP CD8^+^ T cell hyper-proliferation.

### T cell-T cell interactions can promote Itk^−/−^ T cell hyper-proliferation

As CD8^+^ T cells also express MHCI, and T cell-T cell (T-T) interactions have recently been shown to be able to drive the proliferation and effector development of CD8^+^ T cells (26), we examined the role of T-T interaction in lymphopenia-induced CD8^+^ T cell proliferation. In our earlier experiments, we transferred ~120,000 T cells into recipient animals, which could potentially allow T-T interactions in recipients (27). We therefore reduced the number of transferred cells to decrease the frequency or probability of T-T interactions following transfer. WT and *Itk*^−/−^ naïve CD8^+^ T cells were equally mixed at a very low density (2000 cells in total, or 200 *in vivo* assuming 10% take (27)), transferred into *B2m^−/−^Rag^−/−^* recipients and monitored for expansion. Ten days post transfer we found that WT cells hardly proliferated, while *Itk*^−/−^ cells did and formed the predominant HP population (Figure 2G-I). HP of CD8^+^ T cells have been suggested to occur primarily in the relatively smaller compartment of the lymph node, and we indeed saw some WT cells in this compartment when T-T interaction was very limited (Figure 2G, last plot), suggesting that this environment may be able to generate the proximity required for responses by WT cells (28). Furthermore, when we compared the fold expansion over initial numbers of transferred CD8^+^ T cells, we found that when the T-T proximity was reduced, the significantly higher fold expansion of *Itk*^−/−^ cells in *B2m*^−/−^*Rag*^−/−^ recipients was reduced to a similar level to that seen in *Rag*^−/−^ recipients (Figure 2J, *Rag^−/−^B2m^−/−^* recipients with 120,000 or 2000 donor cells). Also notable was that the lack of recipient MHCI allowed a much larger fold expansion (over initial number) by *Itk*^−/−^ CD8^+^ T cells during HP (33-fold expansion by day 10 in the absence of MHCI, compared to 3-fold expansion in the presence of MHCI (co-expansion, Figure 2J, 1^st^ and 2^nd^ blue bars)). This 10-fold difference in expansion suggests a regulatory function for recipient MHCI, presumably via TCR signals, that further tunes the homeostasis of *Itk*^−/−^ HP CD8^+^ T cells. This data suggests that MHCI is required for homeostatic expansion of *Itk*^−/−^ T cells, but that recipient-MHCI acts to modulate this response, and T-T interaction is sufficient to allow this hyper-expansion to occur. These results also suggest that the hyperactive proliferation in the absence of ITK is T cell intrinsic, and these cells are more sensitive to homeostatic signals compared to WT cells, although still regulated by tonic TCR/self-antigen-MHCI interactions.

### ITK signals alter transcriptomic profiles of T cells during HP

We previously reported that innate like memory and HP memory CD8^+^ T cells have distinct transcriptomic profiles (23). While the absence of ITK does not alter the transcriptomic profile of innate memory CD8^+^ T cells (23), we tested whether ITK plays a role in regulating specific gene expression profiles in HP CD8^+^ T cells by collecting WT and *Itk*^−/−^ naïve and HP CD8^+^ T cells for transcriptomic comparison by RNA sequencing. Violin plots shows that global data distribution is similar among all samples (Figure 3A), and principal component analysis (PCA, based on genes with ≥ 2 fold change in any pair of the 4 groups) revealed that *Itk*^−/−^naïve CD8^+^ T cells were similar to WT cells prior to expansion. However, HP for 10 days in *Rag*^−/−^ recipients resulted in significant alteration in *Itkr*^−/−^ CD8^+^ T cell gene expression profile (Figure 3B). Among genes that are significantly altered during HP of WT or *Itk*^−/−^ CD8^+^ T cells, 260 up- and 99 down-regulated genes are common, while a large number of altered genes are singular for WT or *Itkr*^−/−^ cells (Figure 3C). Gene ontology (GO) enrichment analysis further revealed that the genes involved in PCA are over-represented in cellular processes including cell cycle, metabolic regulation, cell communication, immune response to stimuli and other fundamental biological processes (Figure 3D). These data suggest that ITK plays critical function in tuning CD8^+^ T cell HP through a set of key genes that regulates cell proliferation, metabolism and immune cell function, which is specifically triggered by the HP process, as it does not exist in the naïve state.

**Figure 3.**
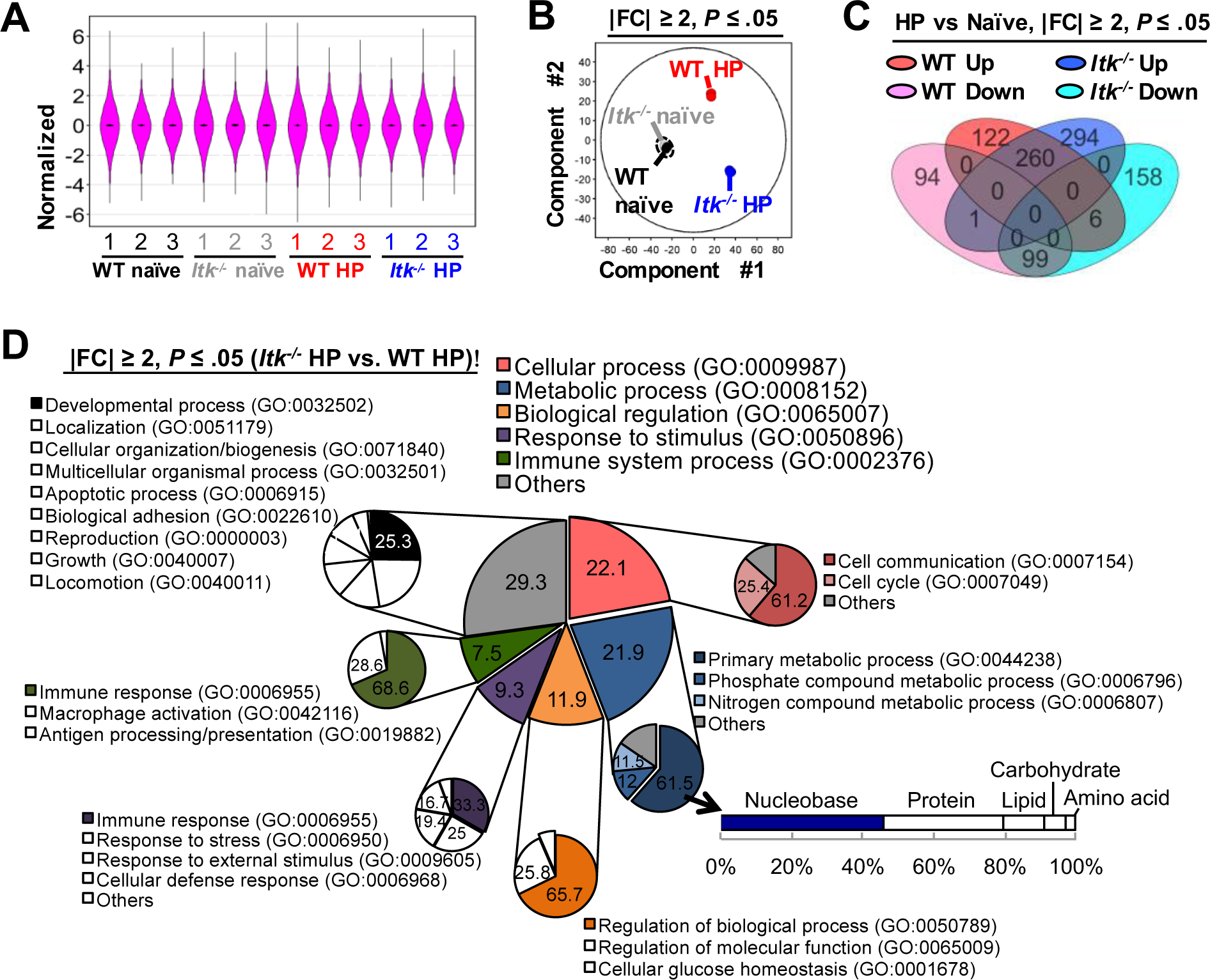
Transcriptomic control by ITK during CD8^+^ T cell homeostatic proliferation under lymphopenic environment. WT and *Itk*^−/−^ naïve and HP CD8^+^ T cells (expanded in *Rag*^−/−^ mice for 10 days) were used for RNA sequencing and analysis. (A) Violin plots of all gene reads of all samples showing similar data quality. (B) Principal component analysis of genes that shown with ≥ 2-fold change in any pairs of group combinations, with FDR (P) ≤ 0.05. Dashed area indicates the error ellipse of all naïve T cell sample profile. (C) Venn diagram of genes that are ≥ 2-fold up- or down-regulated due to the process of HP, with FDR (P) ≤ 0.05. (D) Gene oncology annotation of classified events associated with genes that are with ≥ 2-fold change due to the lack of ITK expression during CD8^+^ T cell homeostatic expansion under lymphopenic condition, with FDR (P) ≤ 0.05. N = 3; FDR: False Discovery Rate.

### ITK regulates T cell proliferation and apoptosis during HP

Gene set enrichment analysis (GSEA) further revealed that genes involved in cell cycle are significantly more enriched in *Itk*^−/−^ HP CD8^+^ T cells compared to WT HP cells 10 days post transfer into lymphopenic recipients, accompanied by the enriched gene set involved in apoptosis (Figure 4A). The rate of cell expansion under these conditions is the integrative result of cell proliferation and death. And we indeed observed that naïve *Itk*^−/−^ CD8^+^ T cells immediately underwent massive expansion, resulting in a > 10-fold increase in the population in 10 days, compared to WT cells (Figure 4B), followed by early collapse of the population, which may be a result of the enhanced apoptotic activity. To better understand the dynamics of CD8^+^ T cell HP and the role of ITK in regulating this process, we established a concise computational model utilizing ordinary differential equations (ODEs) to describe two arbitrary T cell subsets based on their propensity towards proliferating or dying: a proliferating subset (*x*) and a dying subset (*y*), with a conversion index (*c*) that regulates the one-way conversion of cells between these two subsets; the conversion index is determined by a dynamic parameterized combination of the proliferating and dying cell subsets (Figure 4C, & Supplemental Experimental Procedures). Based on random parameter searching optimized through weighted least square regression, we found optimal parameter sets (Figure 4C: best fitting parameters), for the model that recapitulated the actual dynamics observed in experiments (Figure 4B-D). These results suggest that the *Itk*^−/−^ cells have significantly greater potential for proliferation than the WT cells (*k_Itk−/−_* is ~ 9 fold of *k*_WT_) (Figure 4C), as well as a much higher propensity to convert to dying cells over the time period of the expansion, as the conversion index *c* is much larger in *Itk*^−/−^ than in WT cells (Figure 4G). To test the conclusions from these computational simulations, and to provide more insight into the altered dynamics of the CD8^+^ T cell HP by the lack of ITK, we examined the HP CD8^+^ T cells for evidence of proliferation and propensity to undergo apoptosis. The enhanced proliferation of *Itk*^−/−^ CD8^+^ T cells was confirmed by greater dilution of CFSE during the early stages of CD8^+^ T cell HP (Figure 4E). We also analyzed the expression of proliferative (Ki67) and apoptotic (Annexin V) markers of WT and *Itk*^−/−^ HP CD8^+^ T cells at early (day 10) and late (day 75) phases of lymphopenic expansion. We found that *Itk*^−/−^ CD8^+^ T cells exhibited significantly higher signs of proliferation in the early phase (Figure 4F). On the other hand, a significantly higher proportion of *Itk*^−/−^ CD8^+^ T cells also exhibited signs of apoptotic program much earlier than the WT cells (Figure 4H), which is further verified by the observation that *Itk*^−/−^ CD8^+^ T cells significantly up-regulated pro-apoptotic Fas and down-regulated anti-apoptotic Bcl-2 in this process (Figure S2A) compared to WT cells.

**Figure 4.**
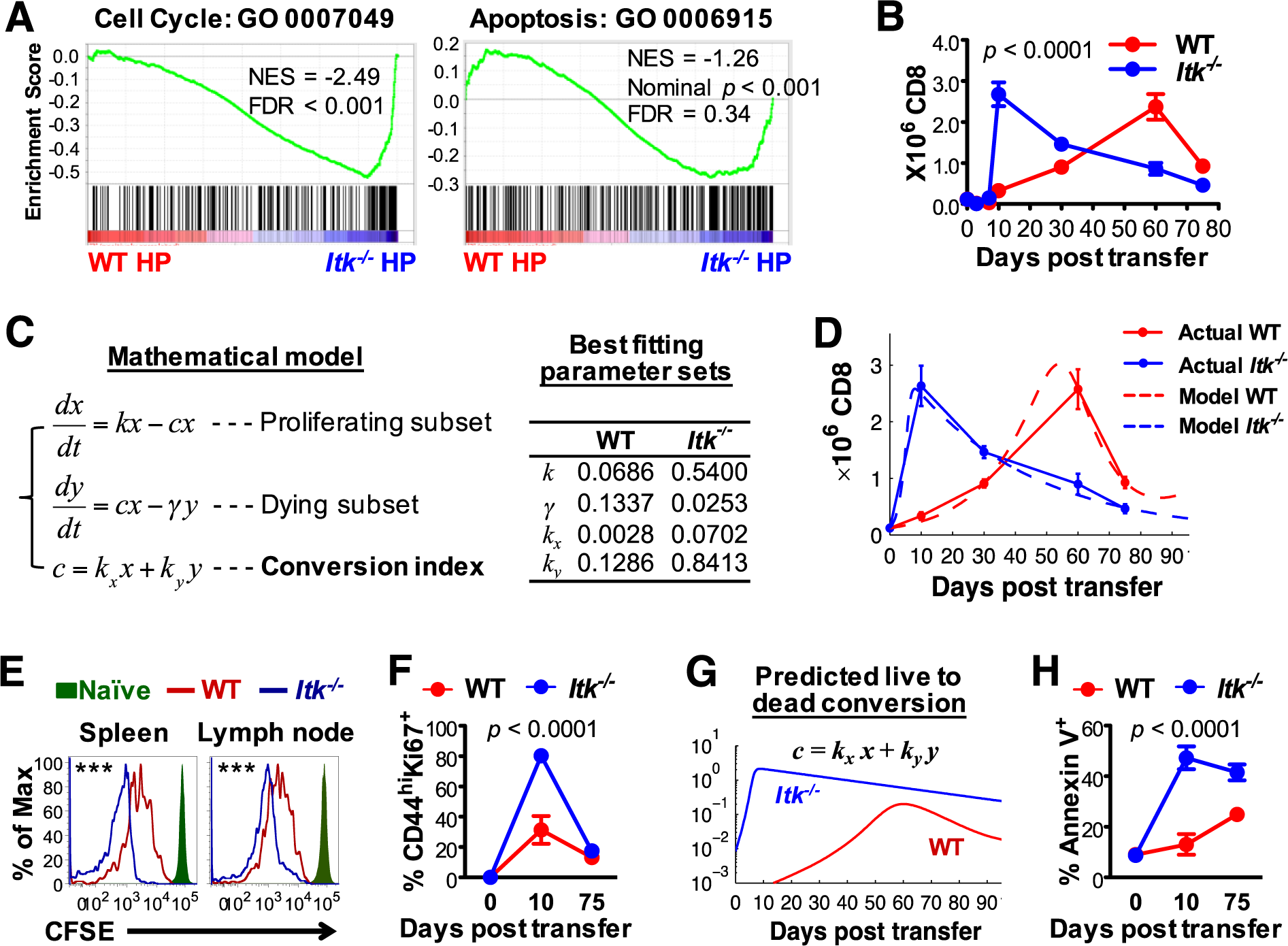
ITK regulates CD8^+^ T cell proliferation and death under lymphopenic environment. (A) Cell cycle and apoptosis-associated gene set enrichment (GO annotated). NES: Net Enrichment Score. (B) Dynamics of WT and *Itk*^−/−^ CD8^+^ T cell HP. 0.12 χ 10^6^ WT or naïve CD8^+^ T cells were adoptively transferred into *Rag*^−/−^ hosts, and CD8^+^ T cells in the spleen of *Rag*^−/−^ recipients at the indicated time points were analyzed. N ≥ 3. Data represent Mean ± SEM; *p* values generated by two-way ANOVA. (C) Ordinary differential equations describing the arbitrary T cell subsets and conversion in **H.L.A.** (**H**uang, **L**uo, and **A**ugust) Model, and best fitting parameter sets as determined by weighted least square regression. (D) Simulated proliferation dynamics recapitulating actual dynamics shown in (B) using the best fitting parameters (overlapping data shown in (B) with that produced by the simulation). (E) Homeostatic proliferation induced division of WT and *Itk*^−/−^ CD8^+^ T cells indicated by the dilution of CFSE. (F) Percentages of proliferative (Ki67^+^) CD8^+^ T cells at initial (day 0), early (day 10) and late (day 75) phase of lymphopenia-induced HP. (G) Predicted live to dead cell conversion by the H.L.A. model using the best fitting parameter sets. (H) Percentages of apoptotic (Annexin V^+^) CD8^+^ T cells at initial, early and late phase of lymphopenia-induced HP.

However interestingly, we noted that the presence of WT cells prevented the collapse of the *Itk*^−/−^ T cell population (first panel in Figure 2F) observed when singly transferred (cf. Figure 4B). When co-expanding with WT congenic T cells, the apoptotic behavior of the *Itk*^−/−^ T cells was similar to WT T cells (Figure S2B, Annexin V^+^), despite significantly enhanced proliferation (Figure S2B, Ki67^+^), suggesting that the observed enhanced apoptosis in these cells upon single transfer (see Figure 4B) is likely cell-extrinsic effect, perhaps due to reduced availability of vital resources per cell under the conditions of the massive expansion of *Itk*^−/−^ cells. It is possible that in the co-transfer model, *Itk*^−/−^ HP T cells compete better for recipient-produced growth/survival resources than WT cells, driving more proliferation of these cells, regulated by ITK in a T cell-intrinsic manner. Alternatively, WT T cells may produce factors for which *Itk*^−/−^ T cells are better able to compete.

### TCR/ITK signals to tune T cell HP and metabolic activity

Lymphopenia-induced T cell HP is extrinsically regulated by common γ-chain (γc) cytokines such as IL-2, IL-7 and IL-15 (29–31). Furthermore, IL-7 induces down-regulation of its receptor IL-7Ra following signaling during HP (32). We found, very interestingly, that the robustly expanding *Itk*^−/−^ CD8^+^ T cells expressed lower levels of IL-2Rβ (the shared subunit of IL-2 and IL-15 receptor complexes) and IL-7Rα, but not the γc or IL-2Rα, compared to the WT cells during lymphopenia-induced HP (Figure S3A&B). The results suggest that *Itkr*^−/−^ T cells may be receiving elevated cytokine signals leading to hyper-proliferation in the lymphopenic environment, and this reduction in receptor expression is likely a result of such signals.

Both the TCR and cytokine signaling pathways are potent in modulating CD8^+^ T cell metabolic activity and we have observed an alteration of the genomic program that regulates the metabolic process in *Itk*^−/−^ HP CD8^+^ T cells compared the WT cells (Figure 3D). In naïve CD8^+^ T cells, the absence of ITK does not affect the basal rate of extracellular acidification (ECAR, an indicator of glycolytic activity) and oxygen consumption (OCR, an indicator of oxidative respiration), but results in increased maximal capacity of ECAR (Figure 5A). Glycolytic metabolism has been reported to support proliferation and effector function of CD8^+^ T cells. To test whether the altered expression of cytokine receptors is related to the cytokine-induced CD8^+^ T cell metabolic activity regulated by ITK, naïve WT and *Itk*^−/−^ T cells were cultured in IL-2, IL- 15 or IL-7 for 12 hours and subjected to ECAR and OCR measurement. We found that *Itkr*^−/−^ CD8^+^ T cells exhibited lower or similar glycolytic (ECAR) and oxidative (OCR) metabolism to WT cells when cultured in IL-2 or IL-15 (Figure S3C&D), however, significantly higher levels of ECAR and OCR in IL-7 (Figure 5B). Intermittent IL-7 signaling activation is intrinsically regulated by TCR signaling and critical for naïve CD8^+^ T cell homeostasis, which can be mimicked by manually switching the culture of cells in media with or without IL-7, or continuously with IL7 (33). By manual mimicry of intermittent IL-7 activation, we found that IL-7-mediated metabolic activities in *Itkr*^−/−^ CD8^+^ T cells was reduced to levels comparable in WT cells (Figure S3F), suggesting IL-7 signaling may be hyper-active in the absence of ITK.

**Figure 5.**
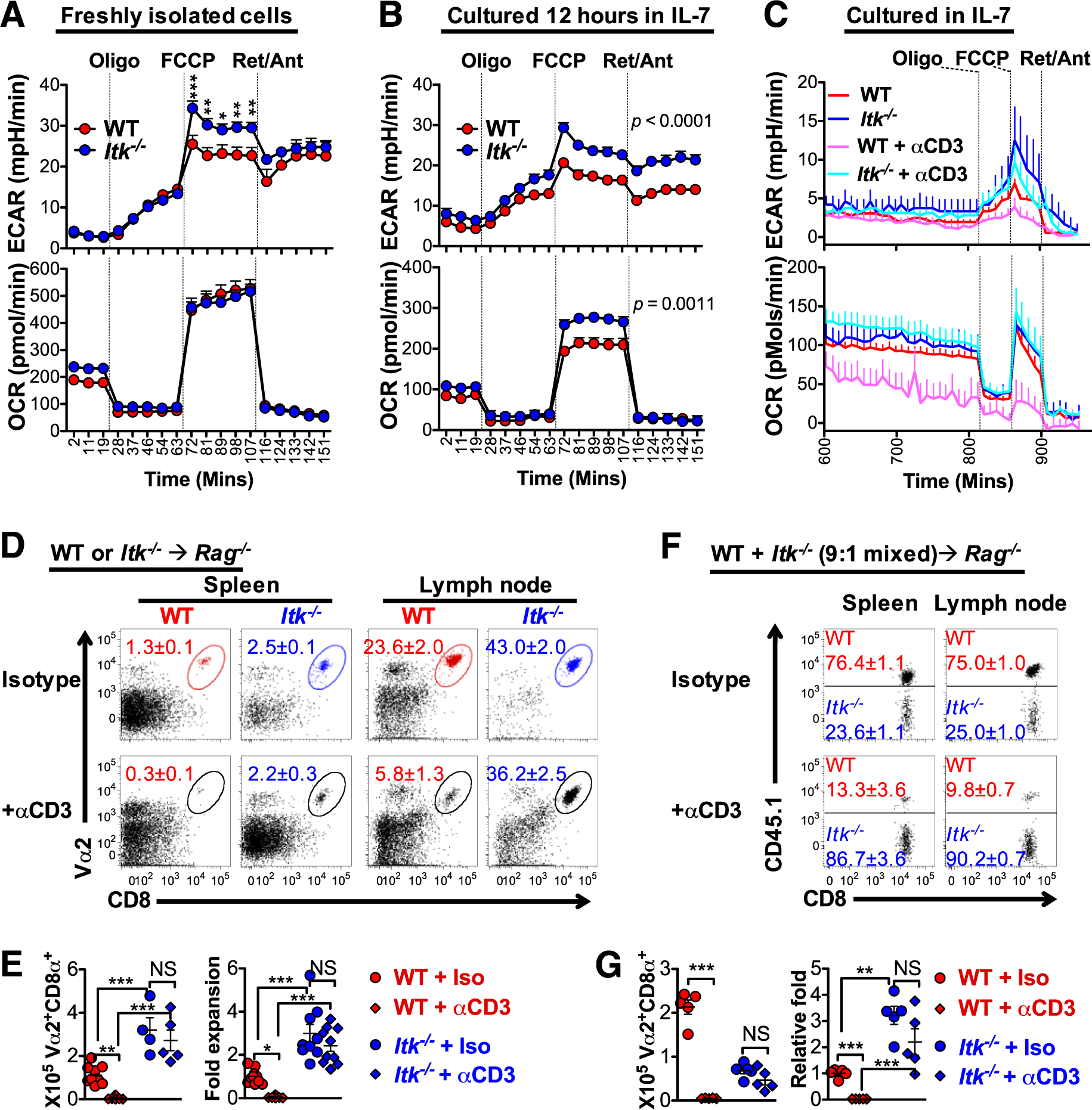
TCR activation tunes down CD8^+^ T cell metabolism and HP *via* ITK signaling. (A) WT or *Itk*^−/−^ naïve CD8^+^ T cells were isolated and immediately analyzed in Seahorse assay media for ECAR and OCR. N = 6. *p* ≤ *0.05, **0.01, ***<0.001 by two way ANOVA with Tukey post test. Data represent results of more than 3 experiments. (B) WT or *Itk*^−/−^ naïve CD8^+^ T cells were cultured in Seahorse assay media with IL-7 overnight, and analyzed for ECAR and OCR. N = 9 pooled from 3 independent experiments. *p* values by two-way ANOVA. (C) WT or *Itk*^−/−^ naïve CD8^+^ T cells were cultured in Seahorse assay media with IL-7 in the presence or absence of αCD3 (0.01 μg/ml) from time 0, and monitored for ECAR and OCR for an overnight. N = 5. (D&E) WT or *Itk*^−/−^ naïve CD8^+^ T cells were transferred into *Rag*^−/−^ recipients on day 0, and recipients received intraperitoneal injection of 0.1 μg/mouse anti-CD3s (or isotype as control) on day 1 and day 3. Spleen and lymph nodes were analyzed on day 10. N = 5. (D) Representative plots of CD8^+^ T cells in the spleen and lymph node on day. (E) Total numbers of donor CD8^+^ T cells in the spleen and lymph node on day 10 and fold expansion by normalization to the number in “WT ^+^ Iso” group. *p* values generated by one-way ANOVA with Tukey post test. (F&G) WT (CD45.1) and *Itk*^−/−^ (CD45.2) naïve CD8^+^ T cells were mixed in a 9:1 ratio (total each mouse) and transferred into *Rag*^−/−^ recipients on day 0, and recipients received intraperitoneal injection of 0.1 μg/mouse anti-CD3s (or isotype as control) on day 1 and day 3. Spleen and lymph nodes were analyzed on day 10. (F) Representative plots of CD8^+^ T cells in the spleen and lymph node on day 10. Cells were pregated on Vα2^+^CD8^+^ and analyzed for CD45.1 to reveal the composition of the final population. (G) Total numbers of donor CD8^+^ T cells in the spleen and lymph node on day 10 and relative fold expansion by normalization to the number in “WT + Iso” group and to the “9:1” ratio of initial population. *p* values generated by one-way ANOVA with Tukey post test.

Triggering the TCR using anti-CD3s antibodies, has been shown to stimulate T cell activation and development *in vivo* (34), and we recently demonstrated that anti-CD3? antibody activates TCR and signals through ITK to drive type 1 regulatory T cell development in multiple organs (35). Given the observation that the lack of ITK, a TCR downstream mediator, is a negative tuner of CD8^+^ T cell HP, we examined the effect of TCR activation in regulating IL-7 mediated CD8^+^ T cell metabolism (Figure 5C and Figure S3G&F). CD8^+^ T cells cultured with anti-CD3s antibody exhibited significantly lower levels of ECAR and OCR that those cultured in IL-7 (Figure S3E). Addition of anti-CD3s antibody at low concentration (0.01 μg/ml) decreased IL-7-mediated ECAR and OCR in WT CD8^+^ T cells, but had no effect in *Itk:*^−/−^ cells (Figure 5C and Figure S3G&H). Our data suggest that TCR activation signals through ITK is a negative tuner of IL-7-mediated metabolic activities in CD8^+^ T cells.

During WT CD8^+^ T cell HP, we observed that administration of low dose of anti-CD3? antibody (0.15 μg/mouse) the day after T cell transfer into *Rag*^−/−^ recipients, resulted in little HP CD8^+^ T cell population (Figure S4A-D). It is unlikely that this is due to depletion of T cells, as administering the antibody 60 days post T cell transfer did not result in reduction in T cell population (Figure S4E&F). However, in the absence of ITK, anti-CD3s antibody did not affect the expansion of *Itk*^−/−^ CD8^+^ T cell HP in *Rag*^−/−^ recipients (Figure 5D&E). A similar effect was observed when mixed WT (initial majority) and *Itk*^−/−^ (initial minority) CD8^+^ T cells were co-transferred into *Rag*^−/−^ mice (Figure 5F&G), suggesting that the T cell-intrinsic TCR signaling through ITK is a negative tuner of CD8^+^ T cells HP under lymphopenic condition.

### ITK signals tune mTOR to regulate T cell expansion and function during HP

CD8^+^ T cells exhibit distinct transcriptional signatures between the short-term effector and long-term memory stages following stimulation (36). GSEA revealed that WT HP CD8^+^ T cells exhibited enriched gene signatures that are found to be up-regulated in long-term memory CD8^+^ T cells (Figure 6A, left), while *Itk*^−/−^HP CD8^+^ T cells had those up-regulated in the effector cells enriched (Figure 6A, right). CD8^+^ T cells undergoing lymphopenia-induced proliferation progressively acquire a memory phenotype, display high levels of CD44 and CD122, however, antigen-inducible early activation and effector markers are typically not expressed (9), and CD62L is not shed, indicative of a lack of effector program (10, 11, 37). However, we found that *Itk*^−/−^ CD8^+^ HP T cells acquire an effector phenotype characterized by CD44^hi^CD62L^lQ^ (38), starting very early during HP as cells were in the significant proliferative phase (Figure 6B), which persisted throughout (Figure 6C&D). This effector phenotype is accompanied by higher expression of CD44, KLRG1 and NKG2D in *Itk*^−/−^ HP cells (Figure 6E). NKG2D-mediated signals are known to augment CD4-unhelped CD8^+^ T cell memory response with enhanced cytotoxicity (39). When stimulated with PMA/Ionomycin, a slightly higher proportion of *Itk*^−/−^ HP CD8^+^ T cells produced IFN-γ and TNF-α, and *Itk*^−/−^ HP CD8^+^ T cells produced more cytokine per cell as indicated by higher mean fluorescence intensity (Figure 6E). This suggests that ITK-mediated signals constrain effector cell fate programing during T cell HP, and removal of such signals allows the development of this effector phenotype.

**Figure 6.**
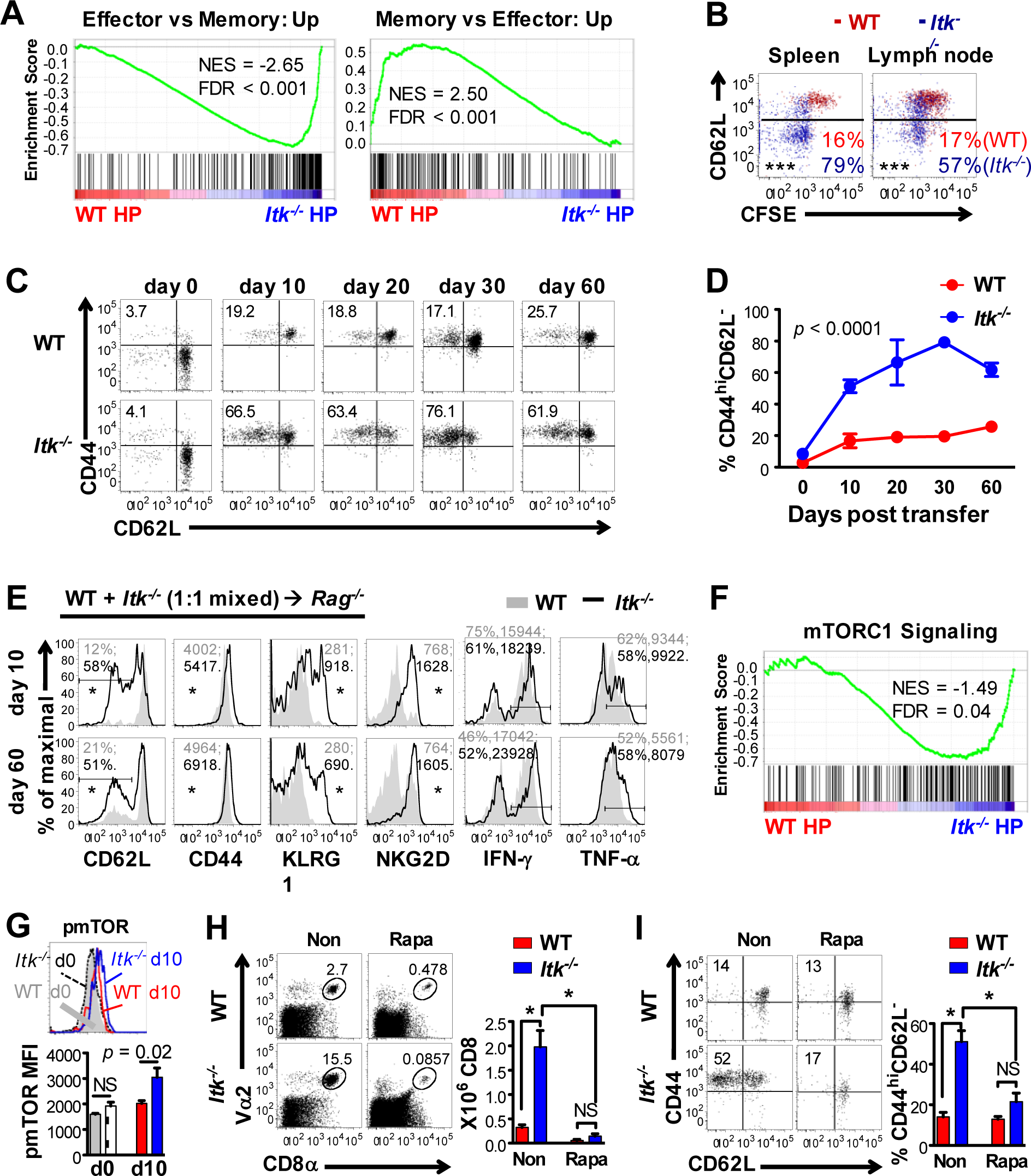
ITK deficiency promotes CD8^+^ T cell lymphopenia-induced expansion and effector programing in an mTOR-dependent manner. (A) Gene set enrichment analysis of genes that are highly enriched in effector versus long-term memory CD8^+^ T cells. NES: Net Enrichment Score; FDR: False Discovery Rate. (B) Representative plots of CFSE dilution and CD62L expression by WT and *Itk*^−/−^ CD8^+^ T cells proliferated in *Rag*^−/−^ lymphopenic hosts for 10 days. Numbers indicate means of 5 replicates, ***p < 0.001 by non-parametric Mann-Whitney test. (C) Representative plots and (D) summary of CD44^hi^CD62L^lQ^ effector phenotype CD8^+^ T cells derived from naïve donor CD8^+^ T cells in the spleen of the lymphopenic *Rag*^−/−^ recipients. Splenic WT and *Itk*^−/−^ OTI-Rag^−−^ CD8^+^ T cells were used as controls on day 0. N ≥ 10 on day 10, and N ≥ 3 at each of the other time points. Data represent Mean ± SEM. *p* value generated by two-way ANOVA. (E) Representative plots of indicated markers, and P/I-induced IFN-γ and TNF-α expression, by CD8^+^ T cells in early (day 10) and late (day 60) phase of lymphopenic proliferation. Numbers on plots show average MFI of ≥ 3 replicates: WT in grey and *Itk*^−/−^ in black; numbers for IFN-γ and TNF-α are for the gated area. *p ≤ 0.05 by Student’s *t* test. (F) Gene set enrichment analysis of mTORCl signaling components. NES: Net Enrichment Score; FDR: False Discovery Rate. (G) Reprehensive plots of phospho-mTOR (pmTOR) of naïve (day 0) and HP (day 10) WT and *Itk-* ^−^ CD8^+^ T cells and summary of pmTOR MFI. N = 3 - 4. (H&I) *Rag*^−/−^ recipients of naïve WT or *Itk*^−/−^ OTI-*Rag*^−/−^ CD8^+^ T cells were treated for 10 days with PBS (Non) or Rapamycin (Rapa). (H) Representative plots and number (total in spleen and lymph node) of WT and *Itk*^−/−^ CD8^+^ T cells expanded 10 days post transfer. (I) Representative plots and percentage of CD44^hi^CD62L^lQ^ effector phenotype CD8^+^ T cells. *p < 0.05, NS = “No Significance”, by non-parametric Mann-Whitney test. N = 5 - 14 from 3 independent experiments. Data represent Mean ± SEM.

During antigen induced CD8^+^ T cell differentiation, T-bet is induced *via* mTOR activation and is essential for effector function, including expression of IFN-γ and related cytotoxic activity (38). By contrast, Eomes expression as a result of low mTOR activity contributes to memory differentiation (see review (40)). We found that *Itk*^−/−^CD8^+^ T cells undergoing HP up regulated T-bet at the early stages (Figure S5B, first panel), with little change in Eomes expression (not shown), compared to WT counterparts. Decreased CD62L and increased T-bet expression along with decreased expression of Foxo3 and Bim are hallmarks of increased mTOR activity (see reviews (40, 41), Figure S5A). In addition, the hallmark transcriptome signatures that are up-regulated in mTORC1 activation are significantly enriched in *Itk*^−/−^ HP CD8^+^ T cells (Figure 6F), and the level of mTOR phosphorylation is significantly increased *Itk*^−/−^ HP CD8^+^ T cells (Figure 6G). Moreover, rapamycin, which targets mTOR, reduced IL-7-mediated ECAR and OCR of *Itk*^−/−^ T cells to the levels observed in WT cells (Figure S3I). These data suggest the possibility that mTOR is hyperactive in the absence of ITK during HP. To examine this, we blocked mTOR activation using rapamycin during the early phase of CD8^+^ T cell HP (over 10 consecutive days), and observed significantly reduced proliferation of *Itk*^−/−^ CD8^+^ T cells to levels seen in WT cells (Figure 6H). Rapamycin also significantly restored T-bet and Bim expression in *Itk*^−/−^ HP CD8^+^ T cells to WT levels (Figure S5C). Most strikingly, blocking mTOR prevented the down-regulation of CD62L in *Itk*^−/−^ HP CD8^+^ T cells undergoing lymphopenic expansion (Figure 6I). These data suggest that mTOR activity is required for the hyperactive proliferation and generation of the effector phenotype in *Itk*^−/−^ CD8^+^ T cells undergoing lymphopenia-induced HP.

### ITK suppresses anti-tumor immunity of HP T cells

WT HP CD8^+^ cells can develop a protective cytotoxic memory-like function indicated by their potent ability to inhibit bacterial and tumor growth in an antigen-specific manner, however, the acquisition of such protective function requires the help of CD4^+^ T cells during the HP process (12). During HP in the absence of CD4^+^ T cells in *Rag*^−/−^ hosts, *Itk*^−/−^ HP CD8^+^ T cells take on an effector-like phenotype (see Figure 6), suggesting that *Itk*^−/−^ HP CD8^+^ T cells might exhibit cytotoxic effector function. To test this, we implanted WT mice (CD45.1^+^) subcutaneously with the T cell lymphoma EL4, or EG7 (an EL4 variant expressing ovalbumin: EL4-OVA, recognized by OT1 T cells), and 2 days post tumor implantation, separately transferred equal numbers of WT or *Itk*^−/−^ HP CD8^+^ T cells (CD45.2^+^) that had been singly expanded in *Rag*^−/−^ recipients. The HP derived cells had no effect in suppressing EL4 growth when the tumor does not express the cognate antigen (OVA) (Figure 7A). Furthermore, in recipients implanted with the EG7 lymphoma, consistent to previous reports (12), WT HP CD8^+^ T cells expanded in the absence of CD4^+^ T cells did not affect tumor growth. However, *Itk*^−/−^ HP CD8^+^ T cells expanded in the absence of CD4^+^ T cells suppressed tumor size by ~ 50% (Figure 7B&C). Although there was a trend towards increased numbers of *Itk*^−/−^ HP CD8^+^ cells at the tumor site, which correlated with shrinkage of the tumors (Figure 7D), we did not observe significant expansion of *Itk*^−/−^ HP CD8^+^ T cells in tumor implanted recipients (Figure 7E). Notably, there was no difference in expression of immunosuppressive markers PD-1/CTLA-4, nor cytotoxic degranulation marker CD107a that might be associated with tumor surveillance/killing (Figure 7F), suggesting that immediate cytotoxicity (likely through cytokine production) mediated by transferred antigen-specific HP effector-like cells may contribute to the tumor shrinkage (42). Indeed, when stimulated with OVA peptide, *Itk*^−/−^ HP CD8^+^ T cells from draining lymph nodes of tumor bearing mice readily responded with increased production of IFN-γ and TNF-a, which was superior to those from WT HP CD8^+^ T cells (Figure 7G). Furthermore, *Itk*^−/−^ HP CD8^+^ T cells that expanded in *B2m^−/−^ Rag^−/−^* hosts retained these enhanced antigen specific anti-tumor responses (Figure S1C-F). The acquisition of anti-tumor activity of *Itk*^−/−^ HP CD8^+^ T cells in the absence of CD4+ T cell help suggests that ITK is a CD8^+^ T cell-intrinsic suppressor for the development of effector function during HP. Furthermore, these results suggest that release of limits placed by recipient MHCI may be a strategy to rapidly enrich the tumor antigen specific effector cells.

**Figure 7.**
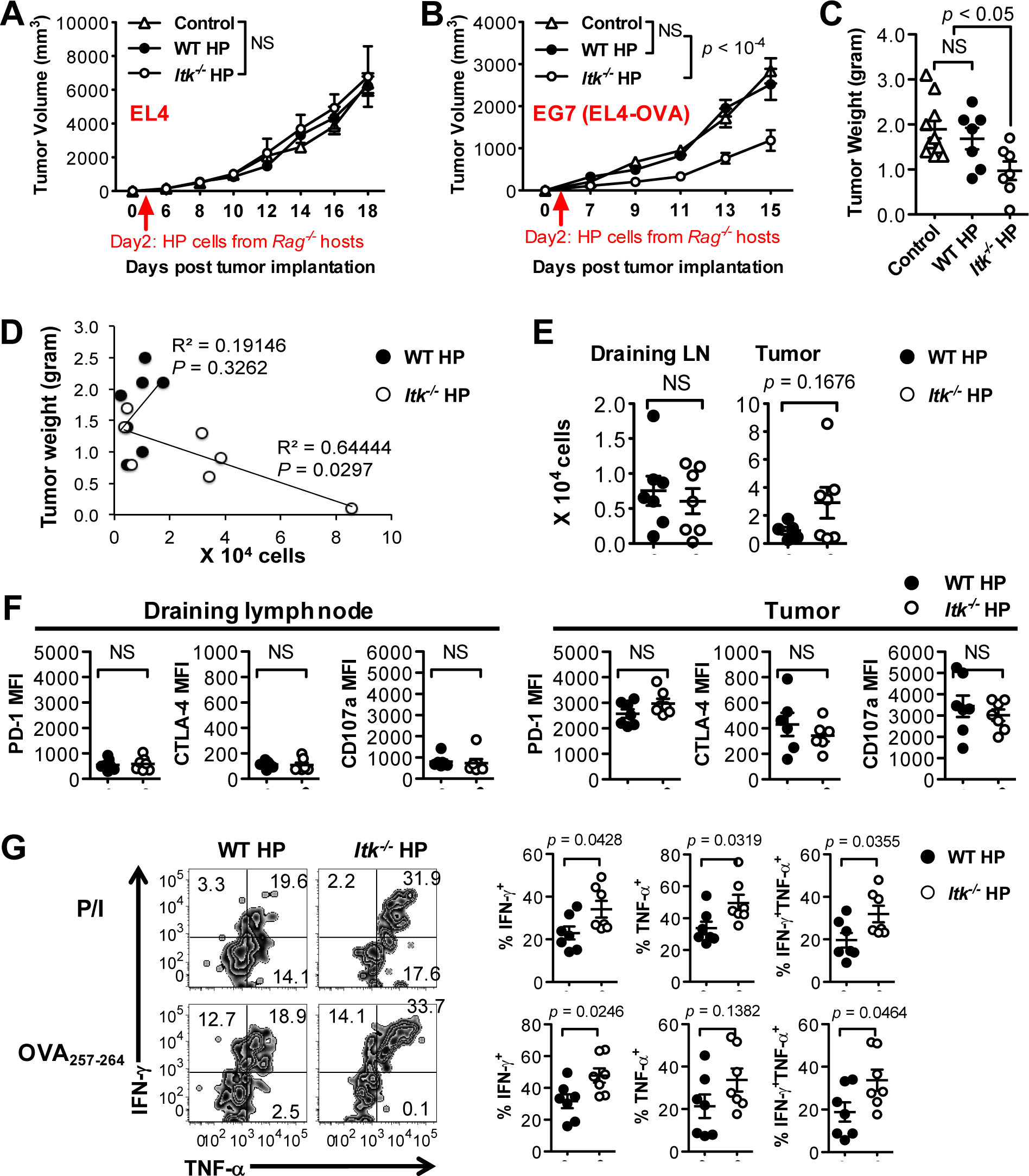
ITK suppresses CD8^+^ T cell anti-tumor immunity developed during lymphopenia-induced HP. CD45.1^+^ WT mice were implanted with EL4 or EG7 (EL4-OVA) lymphoma cells, and received CD45.2^+^ WT or *Itk*^−/−^ HP CD8^+^ T cells (120,000) 2 days later. Tumor size in mice that received no cells (control), WT HP or *Itk*^−/−^ HP CD8^+^ T cells (expanded in *Rag*^−/−^ hosts) along the time course post tumor inoculation. (A) EL4 (N = 7) and (B) EG7 (N = 10) tumor size over time. (C) EG7 tumor weight on day 15. (D-G) CD45.2^+^ HP CD8^+^ T cells in mice shown in (B&C) were analyzed on day 15. (D) Inverse correlation between the number of *Itk*^−/−^ HP CD8^+^ T cells in the tumor site and tumor size. R^2^ implicates the degree of correlation and *P* reflects the likelihood of incorrect prediction. (E) Numbers of CD45.2^+^ HP CD8^+^ T cells in draining lymph node (LN) and tumor site. (F) Expression of PD-1, CTLA-4, and CD107a (MFI) by HP CD8^+^ T cells in draining lymph node and tumor site. (G) CD45.2^+^ HP CD8^+^ T cells from draining lymph nodes of tumor recipients were stimulated with P/I or OVA_257-264_ and IFN-γ and TNF-α production determined by FACS. Data represent Mean ± SEM. *p* values in (A&B) generated by two-way ANOVA, in (C, E, F & G) generated by non-parametric Mann-Whitney test. NS = “No Significance”.

### ITK signals tune cognate antigen/TCR sensitivity of HP T cells

Evaluation of cytokine producing capability of WT and *Itk*^−/−^ T cells following expansion for 10 and 60 days (bypassing the TCR with stimulation with PMA/Ionomycin) revealed a similar or higher percentage of cells secreting IFN-γ and TNF-a in the absence of ITK (Figure 6E). However, when stimulated with OVA peptide, a significantly lower percentage of *Itk*^−/−^ HP T cells responded by secreting IFN-γ during the initial stages of the expansion, but this percentage rose over the time during the expansion, with increased cytokine producing capacity per cell (as indicated by the MFI) (Figure S6A&B, →*Rag*^−/−^ host). This suggests that although *Itk*^−/−^ HP T cells sensed weak TCR signals early during HP with weak cytokine production, they adjusted their sensitivity to TCR signaling during HP. More interestingly, *Itk*^−/−^ HP T cells that had expanded in the absence of lymphopenic recipient MHCI had equal ability to WT cells in producing IFN-γ in response to antigenic stimulation with OVA peptide on day 10, having adjusted their sensitivity to cognate antigen/TCR stimulation at a much early stage compared to when they were transferred into MHCI-expressing recipients (Figure S6A & B, →*B2m^−/−^ Rag^−/−^* host). These data suggest that the absence of host MHCI enhances the tuning of TCR for increased sensitivity in the absence of ITK, which is not observed in WT cells at these antigen concentrations.

Increases in TCR/antigen sensitivity during HP has been suggested to be due to increases in expression of CD8, driven by signals from homeostatic cytokines (43, 44), but we did not observe any significant differences in CD8 expression or TCR expression that could explain the enhanced sensitivity to antigen in the *Itk*^−/−^ HP T cells (Figure S6C). Notably however, this adjustment in *Itk*^−/−^ HP cells was accompanied by changes in the expression of CD5, which has been suggested to be a regulator of TCR antigen sensitivity (45), and was significantly reduced compared to that in WT cells (Figure S6C).

## Discussion

Despite suggestions that there is a positive correlation between TCR signal strength and expansion capability during homeostatic expansion of CD8^+^ T cells under lymphopenic environment (16), we show here that ITK, which positively regulates TCR signal strength, suppresses lymphopenia-induced homeostatic expansion of CD8^+^ T cells. We further show that this occurs in a T cell-intrinsic manner, likely by controlling tonic TCR signaling-mediated tuning of sensitivity to homeostatic cytokines, and fine-tuning of TCR sensitivity to cognate antigen. ITK thus regulates TCR-mediated tuning of CD8^+^ T cell response to homeostatic cytokines and TCR antigen sensitivity for the development of effector phenotype cells and antitumor immunity during HP.

Naïve CD8^+^ T cells undergo HP in response to tonic interactions with self-antigen-MHC and cytokines under lymphopenic conditions such as those due to chemotherapy, irradiation or viral infection (6, 7, 9). These cells expand and develop phenotypic characteristics of authentic antigen-induced memory (37). We found significantly enhanced expansion of *Itk*^−/−^ CD8^+^ T cells in a lymphopenic environment leading to enhanced acquisition of effector phenotypes in a mTOR activity dependent manner. Both TCR and the cytokines responsible for T cell homeostasis can activate mTOR (40), and so whether this dysregulation of mTOR in the *Itk*^−/−^ HP CD8^+^ T cells is downstream of the TCR or cytokines is unclear. We have consistently observed down-regulation of components of the IL-2, IL-15 and IL-7 receptors from the early phase of lymphopenia-induced CD8^+^ T cell HP in the absence of ITK. This may be the result of early cytokine signaling leading to elevated mTOR activity and proliferative responses. Given the enhanced metabolic activities and early expansion observed in the absence of ITK, the dependence on MHCI availability, and the negative effect of TCR activation, we suggest that ITK regulates a signaling pathway downstream of the TCR, which tunes the sensitivity to cytokines that drive the expansion. The resultant elevated cytokine signals may drive elevated mTOR activity leading to the observed responses.

The proliferation of *Itk*^−/−^ T cells is even higher than apparent, since our computational simulations suggest that this is tempered by enhanced death of these cells. It has recently been shown that TCR-activated T cells can generate a soluble version of the extracellular domain of γc c), which can block IL-2R/IL-7R signaling, inhibiting naïve T cell survival (46). However, it is unlikely that this explains the differential survival between WT and *Itk*^−/−^ cells since sγc has very limited effect on the cell population size during homeostasis; furthermore, while the abundance of sγc is positively correlated with γc surface expression (46), we did not see any change in expression of γc on *Itk*^−/−^ T cells. Intermittent tonic TCR signals have been shown to modulate the response of T cells to IL-7 during homeostasis (33). It is therefore more likely that ITK is part of a TCR pathway that suppresses the response of expanding T cells to the IL-7 (and/or other γc cytokines) during lymphopenia, and that its absence results in enhanced cytokine responses leading to mTOR activity and accompanying functional effects.

In support of this idea, *Itk*^−/−^ T cells exhibit different expansion kinetics and apoptotic behavior when transferred alone compared to when they were transferred with WT T cells. Given that tonic MHCI signals are likely not limiting, *Itk*^−/−^ T cells may compete among themselves, likely for cytokine signals, driving massive apoptosis and eventual collapse of the population. This is contrasted with the kinetics when *Itk*^−/−^ T cells were co-transferred along with WT T cells, and it is possible that the co-transferred WT cells provide some resources that alter the kinetics of *Itk*^−/−^ cells. Nevertheless, these data support the view that ITK suppresses the homeostatic response by tuning TCR regulation of cytokine signals as has previously been suggested for naïve T cell homeostasis (33). It has been proposed that expression of MHCI on recipients (or on recipient professional antigen presenting cell (APC)) is a prerequisite for lymphopenia-induced proliferation (9, 47). However our data suggest that the TCR (acting via ITK) and recipient MHCI actually acts as part of a regulatory system that modulates the HP response. Furthermore, the finding that *Itk*^−/−^ T cells expanded 10 fold more in the absence of recipient MHCI supports this idea; i.e. under conditions of limiting MHCI tonic signals, there is less MHCI/TCR tonic feedback acting to inhibit cytokine signals.

In antigen-induced memory differentiation, CD8^+^ T cell-T cell interaction can play a major role in the development of protective function, beyond the role of conventional APC, dependent on adhesion (via LFA-1) or propability of interaction (26). However, the observed superior expansion of small numbers of *Itk*^−/−^ CD8^+^ T cells under conditions of the absence of recipient MHCI is not likely due to alterations in LFA-1 expression (Figure S2C&D, CD11a/CD18). Our results suggest that when TCR signal strength is reduced, tonic MHCI/TCR signals provided by T-T interaction, can support HP.

*Itk*^−/−^ T cells also exhibited accelerated tuning of TCR sensitivity for cognate antigen over time during HP. More importantly, this feature was accelerated when no professional APCs but only T cells expressed MHCI, suggesting that this is negatively regulated by MHCI availability and linked to proliferation. This is remarkable given that the *Itk*^−/−^ CD8^+^ T cells start out with a deficit in that they initially receive weaker TCR signals. The groups of Jameson and Singer have made similar observations of tuning TCR sensitivity in WT CD8^+^ T cells, and suggest that this is a result of altered CD8 expression, which fine-tunes TCR signaling strength over time (43, 44). However, we did not observe any changes in CD8 or TCR expression in the *Itk*^−/−^ CD8^+^ T cells over this period. Instead we found that CD5 expression was lower on the *Itk*^−/−^ CD8^+^ T cells over time compared to WT cells, suggesting that CD5 expression may regulate this tuning as well. Alternatively, this tuning may occur by alterations in intracellular signaling molecules that were not examined. Nevertheless, these data support the conclusion that TCR and homeostatic cytokines can tune the sensitivity of subsequent antigenic responses, and our data using the *B2m^−/−^ Rag^−/−^* mice, where this is even more enhanced, support the view that strong and/or frequent TCR signals triggered by tonic MHCI interaction limit this enhanced tuning of the response.

One significant function of HP CD8^+^ T cells is their anti-tumor activity, which may benefit tumor therapy following lymphopenia created by chemo- or radio-therapy (13, 14). Lymphopenia-induced generation of anti-tumor CD8^+^ T cells require the presence of CD4^+^ T cells (12), which creates a dillema of having both CD8^+^ T cell anti-tumor immunity, and potential CD4^+^ T cell autoimmunity. Our findings that the absence of ITK in CD8^+^ T cells during lymphopenia-induced expansion gives rise to significant anti-tumor protection are promising in this regard in that it potentially allows the development of CD8^+^ T cell anti-tumor immunity in the absence of CD4^+^ T cells.

Taken all together, our data support a model where the TCR, acting *via* ITK, tunes the response of naïve CD8^+^ T cell homeostatic expansion, effector function development, and the sensitivity of subsequent antigenic responses, which suggest that strong and/or frequent TCR signals limit these responses. This model also suggests an approach to generate potent and high sensitivity anti-tumor CD8^+^ T cells for therapy.

## Materials and Methods

### Study design

The aim of this study was to determine the role of TCR signaling *via* non-receptor tyrosine kinase ITK in T cell homeostasis and effector function under lymphopenic conditions. We controlled for T cell antigen receptor/MHC affinity by generating *Itk*^−/−^ (and WT) mice carrying an ovalbumin (OVA) specific T cell antigen receptor (OTI) on a *Rag* deficient, C57BL/6 background, allowing us to focus specifically on TCR signaling via ITK. We isolated naïve CD8^+^ T cells from these mice and compared their homeostatic proliferation, effector cell fate programing, metabolic activities and antigen-specific anti-tumor functions.

### Mice

NSG (NOD.Cg-*Prkdc^scid^Il2rg^tm1W7^*/SzJ, H2k^d^), CD45.1^+^ (*B6.SJL-Ptprc^a^ Pep3^b^*/BoyJ), and *B2m^−/−^ (B6.129P2-B2m^tm1Unc^*) mice were from The Jackson Laboratory (Bar Harbor, ME). OTI *Rag^−/−^ (B6.129S7-Rag1^tm1Mom^* Tg(TcraTcrb)1100Mjb N9+N1) and *Rag^−/−^yc^−/−^* (B10;B6-*Rag2^tm1Fwa^II2rg^tm1Wjl^*) mice were purchased from Taconic (Hudson, NY). *Itk*^−/−^ OTI *Rag*^−/−^ mice were generated by crossing *Itk*^−/−^ and OTI *Rag*^−/−^ mice. CD45.1+ OTI *Rag*^−/−^ mice were generated by crossing CD45.1+ (B6.SJL-Ptprc^a^ Pep3^b^/BoyJ, Jackson) and OTI *Rag*^−/−^ mice. *B2m^−/−^ Rag^−/−^* mice were generated by crossing *B2m^−/−^ (B6.129P2-B2m^tm1Unc^*, Jackson) and *Rag*^−/−^ mice. NSG was on a NOD/ShiLtJ (H2k^d^) background and all other mice were on a C57BL/6 (H2k^b^) background. All experiments were approved by the Office of Research Protection’s Institutional Animal Care and Use Committee at Cornell University.

### Cell purification and adoptive transfer

CD8^+^ naïve T cells were purified from age matching OTI *Rag*^−/−^ (referred to as **WT**) and *Itk*^−/−^ OTI *Rag*^−/−^ (referred to as ***Itk*^−/−^**) littermates through magnetic negative selection. To enrich CD8^+^ naïve T cells: splenocytes were stained by biotin-conjugated antibodies against Gr-1, CD49b, Ly-76, NK1.1, F4/80, CD11c, c-Kit, CD25, CD11b, and CD44, followed by anti-biotin MicroBeads (Miltenyi Biotec, Cambridge, MA). Cells passed through MS column (Miltenyi) were then subjected to flow sorting, sorted cells with purity (CD8α^+^TCRVβ5^+^CD44^lo^CD122-) greater than 95% were used for adoptive transfer into *Rag*^−/−^ recipients, through retro-orbital injection (100,000 ~ 500,000 cell/mouse as specified). To transfer CD8^+^ T cells into sub lethally irradiated recipients, CD45.1^+^ congenic mice were subjected to γ-irradiation (600 cGy), followed by retro-orbital injection of CD45.2^+^ WT or *Itk*^−/−^ naïve CD8^+^ T cells (5 × 10^5^/mouse) 24 hours post irradiation.

### Antibodies and flow cytometric staining

All fluorochrome-conjugated antibodies used are listed in “fluorochrome-target (annotation and clone)” format as follows: eFluor 450-CD122 (IL-2Rβ/IL-15Rα, TM-b1), eFluor 450-H-2Kb (AF6-88.5.5.3), FITC-IL-2 (JES6-5H4), PE-T-bet (eBio4B10), Alexa Fluor 700-CD45.2 (104), PerCP-Cy5.5-CD 127 (IL-7Ra, A7R34), PerCP-eFluor 710-Eomes (Dan11mag), PE-Cy7-IFNγ (XMG1.2), and PE-Cy7-NKG2D (CX5) were purchased from eBioscience (San Diego, CA); V500-CD44 (IM7), FITC-CD45.1 (A20), FITC-CD45.2 (104), FITC-Fas (Jo2), FITC-TCRVβ5 (MR9-4), PE-Annexin V, PE-CD18 (C71/16), PE-IL-4Ra (mIL4R-M1), PE-TCRVa2 (B20.1), PE-TNF-a (MP6-XT22), PE-CF594-CD8a (53-6.7), Cy5-Annexin V, Alexa Fluor 700-CD62L (MEL-14), Alexa Fluor 700-Ki67 (B56) and Allophycocyanin-Cy7-TCRVa2 (B20.1) were purchased from BD Biosciences (San Diego, CA); PE-Texas Red-CD8a (5H10) were purchased from Invitrogen (Carlsbad, CA); FITC-Bcl-2 (10C4), FITC-KLRG1 (2F1), FITC-CD11a (2D7), Allophycocyanin-CD132 (γc, TUGm2), Alexa Fluor 700-CD45.1 (A20), PE-Cy7-CD5 (53-7.3), and PE-Cy7-CD62L (MEL-14) were from Biolegend (San Diego, CA). Biotin-conjugated anti-Gr-1 (RB6-8C5), CD49b (DX5), and Ly-76 (TER119) were from BD Biosciences, anti-NK1.1 (PK136), B220 (RA3-6B2), F4/80 (BM8), CD11c (N418), c-Kit (2B8), CD25 (PC61.5), CD11b (M1/70), and CD4 (GK1.5) were from eBioscience, and anti-CD44 (IM7) was from Biolegend. Cells were resuspended and blocked with anti-mouse CD16/CD32 antibody (Fc block, eBioscience) in 2% fetal bovine serum containing PBS (flow buffer), followed by surface staining by the indicated antibodies. After surface staining: cells subjected to cytokine and nuclear staining were fixed and permeabilized with the Foxp3 staining buffer set (eBioscience), followed by incubation with antibodies; cells subjected to apoptotic assay were stained for Annexin V and PI/RNase as previously described (48). Stained cells were analyzed on LSRII system (BD Biosciences). Data were analyzed using FlowJo software (Tree Star Inc., OR).

### In vitro stimulation

For analysis of IFN-γ and TNF-a production by expanded cells, splenocytes from recipient mice were collected, plated at 2 × 10^6^ cells/ml, and either left unstimulated, or stimulated with 100 ng/ml PMA/0.5 μM Ionomycin (Sigma) (referred to as **P/I**), or 1 μg/ml OVA257-264 peptide (Peptides International, Louisville, KY), in the presence of 5 - 10 μg/ml Brefeldin A (**BFA**, Sigma) for about 5 hours, followed by staining and flow cytometric analysis.

### CFSE cell division assay

5 × 10^6^/ml WT or *Itk*^−/−^ OTI-*Rag*^−/−^ naïve CD8^+^ T cells were incubated with 10 μM CellTrace CFSE dye (Life Technologies) in 37°C for 10 mins., followed by extensive washing. CFSE labeled cells per recipient mouse were then transferred to lymphopenic hosts and analyzed by flow cytometry in the indicated time point.

### Rapamycin treatment

Rapamycin (**Rapa**) was purchased from LC Laboratories (Woburn, MA). 10 mg/kg (Rapa/body weight) was delivered via intraperitoneal daily injection for 10 consecutive days following adoptive transfer.

### Quantitative real time-PCR

TCRVα2^+^CD8α^+^ HP cells were sorted from mock-treated (Non) or rapamycin-treated (Rapa) *Rag*^−/−^ recipients (pooled spleen and lymph nodes) 10 days post transfer and treatment, on a FACSAria Cell Sorter (BD Biosciences). Sorted cells were subjected to RNA isolation using RNeasy Mini Kit (Qiagen, Valencia, CA), followed by cDNA synthesis using You Prime First-Strand Beads Kit (GE Healthcare, Piscataway, NJ). Quantitative real-time PCR (qRT-PCR) was carried out using Taqman probe sets for Foxo3 (Mm01185722_m1) and Bim (Mm00437796_m1) (Life Technologies, Grand Island, NY).

### Anti-CD3s administration in vivo

On the specified dates post T cell transfer into *Rag*^−/−^ mice, recipient mice were injected with 0.1-0.15 μg/mouse anti-CD3ε (145-2C11) intraperitoneally.

### RNA sequencing, gene profiling, and gene set enrichment analysis

Naïve CD8^+^ T cells and HP CD8^+^ T cells (isolated from female origin) expanded 10 days in *Rag*^−/−^ recipients were using for total RNA extraction following by RNA sequencing (Illumina HiSeq 2500) in the RNA Sequencing Core Facility at Cornell University. The processed abundance measurement values were used in GeneSpring GX (Agilent, Santa Clara, CA) for violin plot quality control, principal component analysis (PCA) and gene differential expression. Genes with greater than 2-fold change in *Itk*^−/−^, compared to WT HP CD8^+^ T cells were subjected to Gene Oncology (GO) annotation and classification using the Protein ANalysis THrough Evolutionary Relationships (PANTHER) classification system (49). Gene Set Enrichment Analysis (GSEA) is done using the GSEA platform developed by the Broad Institute (50), and gene set enrichment with false discovery rate (FDR) ≤ 0.05 are considered significant.

### Seahorse metabolic assay

CD8^+^ T cell extracellular acidity rate (ECAR) and oxygen consumption rate (OCR) were detected using the XF24 Extracellular Flux Analyzer (Seahorse Bioscience, MA), in assay medium (pH = 7.35) containing 10 ~ 20 mM glucose, 1mM L-glutamine, 1 mM sodium pyruvate, and 100 U penicillin/streptomycin, with oligomycin (1 μM), phenylhydrazone (FCCP, μM), Rotenone/antinomycin (0.5 μM), and/or 2-deoxy-glucose (2-DG, 100 mM). Naïve CD8^+^ T cells were freshly analyzed or following treatment with IL-2 (20 ng/ml), IL-7 (20 ng/ml) or IL-15 (20 ng/ml), anti-CD3s (high dose at 0.1 μg/ml, low as 0.01 μg/ml), with/or without rapamycin (20 ng/ml) as indicated for overnight. Intermittent IL-7 stimulation was done by switching cells between medium with or without 20 ng/ml IL-7, every 12 hours for 3-5 days.

### Tumor inoculation and HP CD8^+^ T cell treatment

Lymphoma cell lines EL4 and EG7 (EL4 with transgenic expression of ovalbumin: EL4-OVA) were purchased from American Type Culture Collection (Manassas, VA). 0.2-0.5 × 10^6^ tumor cells were injected subcutaneously into the flanks of the backside of CD45.1 mice. 2 days post tumor implantation, HP T cells (120,000 WT or *Itk*^−/−^ OTI *Rag*^−/−^ CD8^+^ T cells, previously expanded for 75 days in *Rag*^−/−^ recipients or 25 days in *B2m*^−/−^*Rag*^−/−^ recipients) were adoptively transferred by retro-orbital injection. Tumor diameters (major and minor) were measured with calipers at indicated time points, and analyzed using an ellipsoid volume formula: Volume = 0.524 x major diameter x minor diameter^2^. At the endpoint of the experiment, tumors were excised and weighed. Tumors and draining lymph nodes (axillary, brachial and/or inguinal lymph nodes on the side of the tumor) were collected to determine the number of transferred HP CD8^+^ T cells. Cells from draining lymph nodes were stimulated with PMA/Ionomycin or OVA257-264 peptide with BFA to determine cytokine production.

### Computational simulation

The H.L.A. model is described in details in the Supplementary information. To solve the ODEs in Figure 4C, random parameter searching was performed through weighted least square regression, using MATLAB (MathWorks, Natick, MA).

### Statistical analysis

Statistical graphs present “Mean ± S.E.M”. Non-parametric Mann-Whitney test, unpaired two-tailed Student’s *t* test, one-way ANOVA and two-way ANOVA with Tukey’s post-hoc test between groups were performed using GraphPad Prism version 5.00 for Macintosh (GraphPad, San Diego, CA). A difference with*p* < 0.05 is considered significant. Linear regression was carried out using Office Excel (Microsoft). “NS” denotes “No Significance”.

## Acknowledgments

We thank Dr. W. Iddings, S. Solouki, L. Zhang, A. Redko, Dr. L. Huang and Dr. Y. Xu for technical help; Drs. J. Appleton, M. Bynoe, and E. Denkers for sharing *Rag*^−/−^ mice; Dr. J. Grenier for help in RNA sequencing and raw data processing. We also thank Drs. M.O. Li at Memorial Sloan Kettering Cancer Center and B.D. Rudd at Cornell University for helpful discussions. Funding: This work was supported by grants from the National Institutes of Health (AI120701, AI138570 and AI126814 to A.A., AI129422 and AI138497 to A.A. and W.H., and AI137822 to W.H., HD076210 to the Cornell RNA Sequencing Core Facility, GM110760 to the Center for Experimental Infectious Disease Research at Louisiana State University), Howard Hughes Medical Institute (HHMI Professorship to A.A.), the American Association of Immunologists (Careers in Immunology Fellowship to W.H.), and Louisiana State University (Faculty Development Award to W.H.). Author contributions: W.H. and A.A. conceived research, designed experiments, analyzed and interpreted data, and wrote the manuscript; W.H. and J.L. perform computational simulation and wrote the methods. Competing interests: AA. has received sponsored research support from 3M. Other authors declare no competing financial interests. Data and materials availability: Raw flow cytometry data is available upon request. The sequencing data were deposited in NCBI GEO with access number GSE72703, and the reviewers’ link is: http://www.ncbi.nlm.nih.gov/geo/query/acc.cgi?token=ihmxscainjkvjib&acc=GSE72703. Source code of the computational simulation is provided in a supplementary zipped folder.

## Supplementary Methods and Materials

**H.L.A. Model: Lymphopenia-induced CD8^+^ T cell homeostatic proliferation.**

Authors W.**H.**, J.**L.**, and A.**A.** contributed to the computational model, thus the model is named after the authors’ last names as “**H.L.A.**”.

In the experiments shown in Figure 4B of the parent manuscript, we observed that the number of CD8^+^ T cells expanded under lymphopenic condition has a non-monotonic tendency along with time, which is dramatically different from the classical logistic model. We therefore tried to build a new mathematical model to fit the experimental observations, and determine possible biological processes involved in the CD8^+^ T cell homeostasis.

First, we proved that a model composed of identical cells would fail to fit the observed non-monotonic dynamics. If we assume that all HP CD8^+^ T cells are identical, the cell number *y* would have a single rate of change:

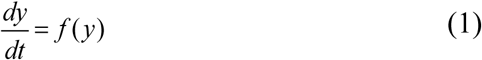

Then logistic model as the example in (2) could be a potential solution:

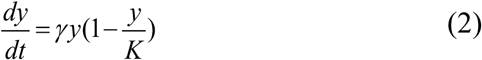

However, the population dynamics observed in our experiments is non-monotonic (see Figure 4B of the parent manuscript). When the *y-t* curve is non-monotonic, one can always find two points **A** and **B** (HLA Figure 1).

**HLA Figure 1.**
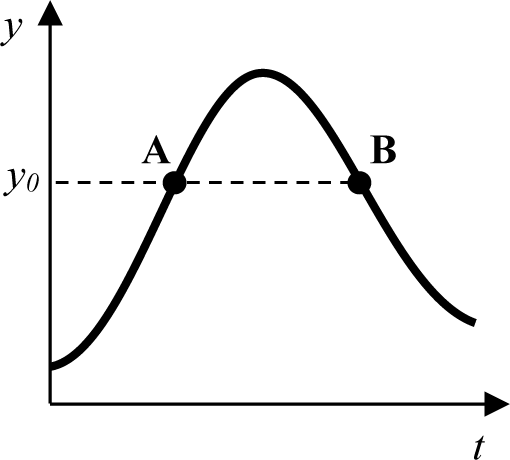
Example of non-monotonic dynamics with *yo* corresponding to distinct dynamics at point A and B.

Both **A** and **B** correspond to the same *y* value (*yo*), but one is in the increasing phase (point **A**) and the other in the decreasing phase (point **B**). i.e.

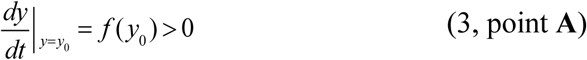

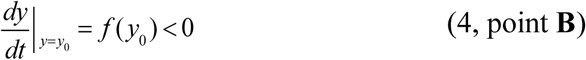

(3) and (4) are contradictory if the dynamics of HP CD8^+^ T cells are contributed by identical cells as shown in (1).

Hence, we designed a two-component model for CD8^+^ T cell homeostatic proliferation under lymphopenic condition. We assume that HP CD8^+^ T cells have two major arbitrary subsets: one with a propensity to continue to proliferate (*x*) as well as convert to the other, which has a propensity towards dying (*y*). The dynamics of these populations are controlled by the proliferation index (*k*), conversion index (*c*), and death index (γ). The changing rate of these two subsets follows the following dynamics:

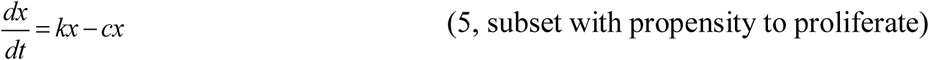

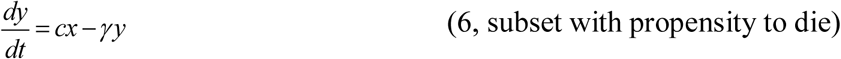

The prolife rating cells replicate themselves with proliferation index *k* and convert to the dying cells with conversion index *c* (equation (5)). The dying cells come from the converted cells and cannot replicate, with a death rate controlled by death index γ (equation (6)). While the proliferating index and dying index are constant, the conversion index that reflects the conversion of proliferating cells to cells with propensity towards dying is influenced by the abundance of both proliferating and dying subsets. We used a weighted parameterization strategy to include these factors in our model:

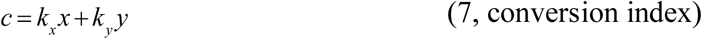

Here, *k*_*x*_ and *k*_*y*_ reflect the influence of proliferating subset and dying subset on the conversion process respectively.

We used the weighted least square regression approach to determine the best fitting parameter sets. In brief, we used MATLAB (MathWorks, Natick, MA) to perform random parameter searching by weighted least square regression to fit the model above (ordinary differential equations [ODE] (5), (6), and (7), as in Figure 4C of the parent manuscript) based on the observed experimental dynamics. The weighted square error is defined by the following equation: 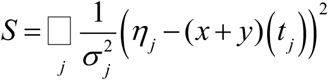, where *η*_*j*_ is the mean of the actual T cell numbers observed in experiment at time point *t*_*j*_, *σ_j_* is the standard error of *η*, and (*x+y)(t*_*j*_) is the total simulated T cell number by the ODE model. The initial value of the parameter optimization covers the biologically meaningful parameter space to avoid *S* falling into a local minimum. For each set of parameters, a corresponding (*x+y)(t_j_)* was compared to the actual experimental data to get its weighted square error *S*. The parameter set is optimized to minimize the weighted square error S, i.e. deviation of simulated dynamics from actual dynamics observed in experiments. Initial CD8^+^ T cells transferred to the lymphopenic recipients belong to proliferating subset, and there are no dying cells at *to* = 0; thus our initial values were set as (*x0, y0*) = (120000, 0).

Fitting commands with detailed information of equation structure, solution, and plotting are available in “HLA_Simulation\main.m”; experimental data input is in “HLA_Simulation\data_file.xlsx”; and results are output in two sections including the parameters in “HLA_Simulation\result_parameters.xlsx” and figures in “HLA_Simulation\result_figures’\ To simulate alternative non-monotonic population dynamics, one can replace population data in “data_file.xlsx”, and seek output in “result_rapameter.xlsx” and “result_figures”.

Based on the actual dynamics observed in our experiments, our results compose two sections including 1) best fitting parameter sets for WT and *Itk*^−/−^ CD8^+^ T cell HP are as presented in Figure 4C of the parent manuscript; and 2) figures comparing simulated and actual dynamics (result_figures\figure_1, as presented in Figure 4D of the parent manuscript), or showing values as annotated in Table S1 and presented in Figure S7.

**Figure S1.**
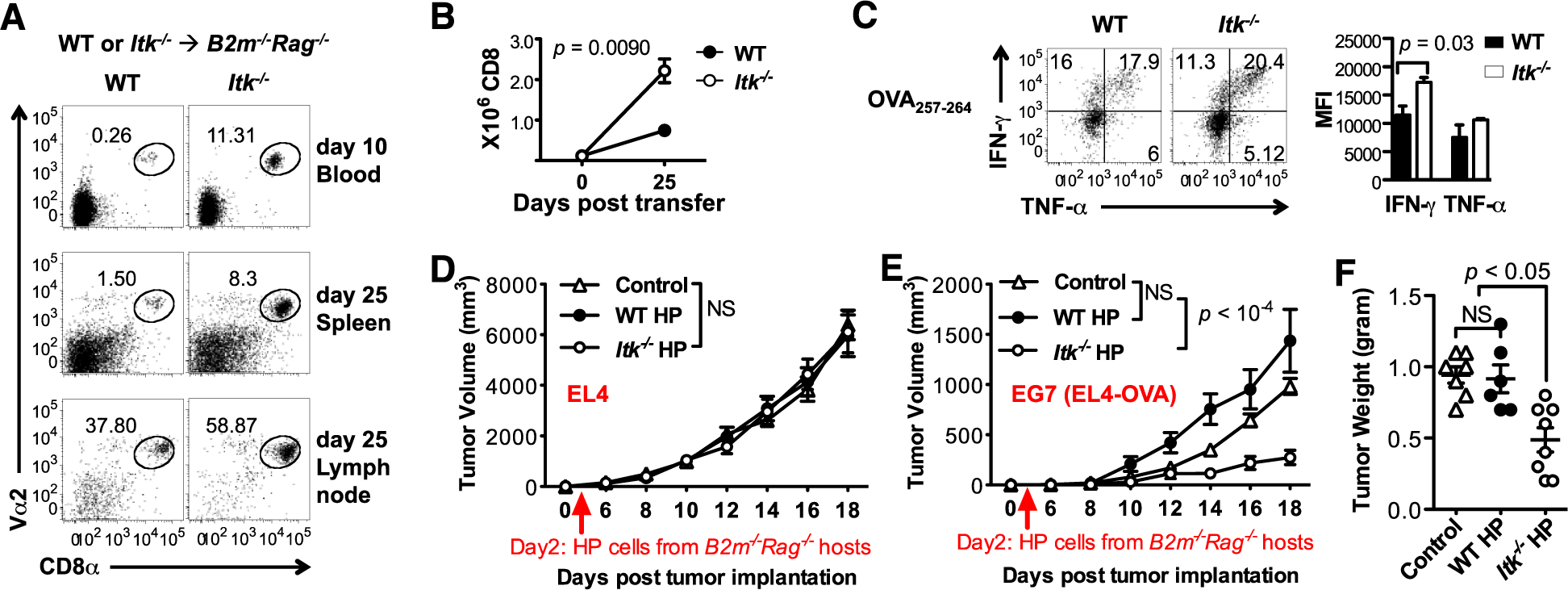
Expression of recipient MHCI is dispensable for ITK-mediated negative tuning of expansion, antigen sensitivity, and anti-tumor immunity of HP CD8^+^ T cells. (A-C) 120,000 naïve WT or *Itk*^−/−^ OTI-*Rag*^−/−^ CD8^+^ T cells are transferred into *B2m^−/−^ Rag^−/−^* recipients. N = 3. (A) Representative plots indicating abundance of WT and *Itk*^−/−^ CD8^+^ T cells expanded 10 days (blood) or 25 days (spleen and lymph node) post transfer. (B) Number (sum of spleen and lymph node) of WT and *Itk*^−/−^ CD8^+^ T cells expanded 25 days post transfer. (C) Representative plots and summary of MFI of IFN-γ and TNF-α expression induced by MHC1/OVA257-264 peptide stimulation of CD8^+^ T cells expanded 25 days in spleen of *B2m*^−/−^*Rag*^−/−^ hosts. (D-F) EL4 or EG7 (EL4-OVA) lymphoma cells were subcutaneously implanted in the flank of mice (CD45.1^+^) in a contralateral manner, followed by infusion of CD45.2^+^ WT or *Itk*^−/−^ HP OTI-*Rag*^−/−^ CD8^+^ T cells (previously expanded for 25 days in *B2m*^−/−^*Rag*^−/−^ hosts as in A-C) 2 days later. N = 7 - 9. (D) EL4 and (E) EG7 tumor size in mice that received no cells (control), WT HP or *Itk*^−/−^ HP CD8^+^ T cells along the time course post tumor inoculation. (F) EG7 tumor weight on day 18. Data represent Mean ± SEM. *p* values in (B, D & E) were generated by two-way ANOVA, in (C & F) were generated by Student’s *t* test. NS = “No Significance”.

**Figure S2.**
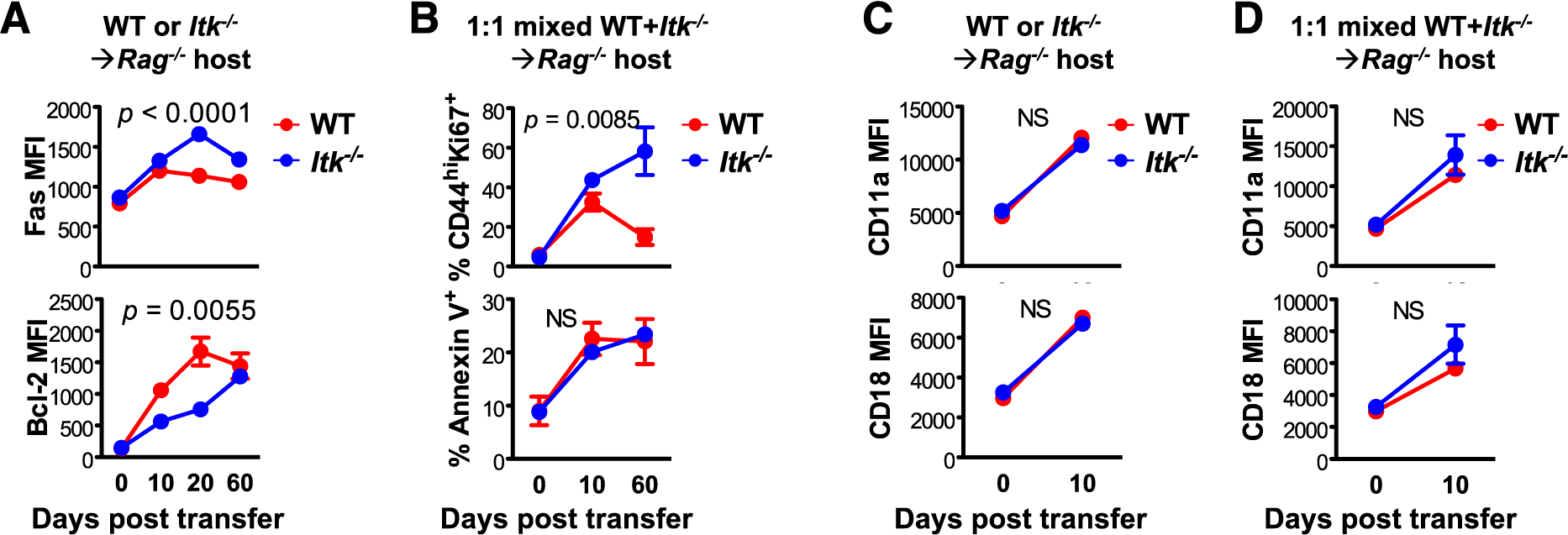
Dynamic of expression of apoptotic, proliferative, survival and migratory markers by the HP CD8^+^ T cells. (A&C) 120,000 naïve WT or *Itk*^−/−^ OTI-*Rag*^−/−^ CD8^+^ T cells are transferred into *Rag*^−/−^ recipients. N = 3–7. (B&D) Congenic naïve WT (CD45.1^+^) and *Itk*^−/−^ (CD45.1^−^) OTI-*Rag*^−/−^ CD8^+^ T cells were mixed at a ratio of 1:1, and a total 120,000 T cells were transferred to *Rag*^−/−^ recipients. N = 3. (A) Dynamics of expression (Mean Fluorescent Intensity, MFI) of Fas and Bcl-2. (B) Percentage of CD44^hi^KI67^+^ and AnnexinV^+^ cells over CD8^+^ T cells expanded during lymphopenia-driven HP in the co-transfer model. (C&D) Dynamics of expression (MFI) of LFA subunits CD11a and CD18 in the singly transferred (C) and co-transferred (D) cells. Data represent Mean ± SEM. *p* values were generated by two-way ANOVA. Data represent Mean ± SEM. NS = “No Significance”.

**Figure S3.**
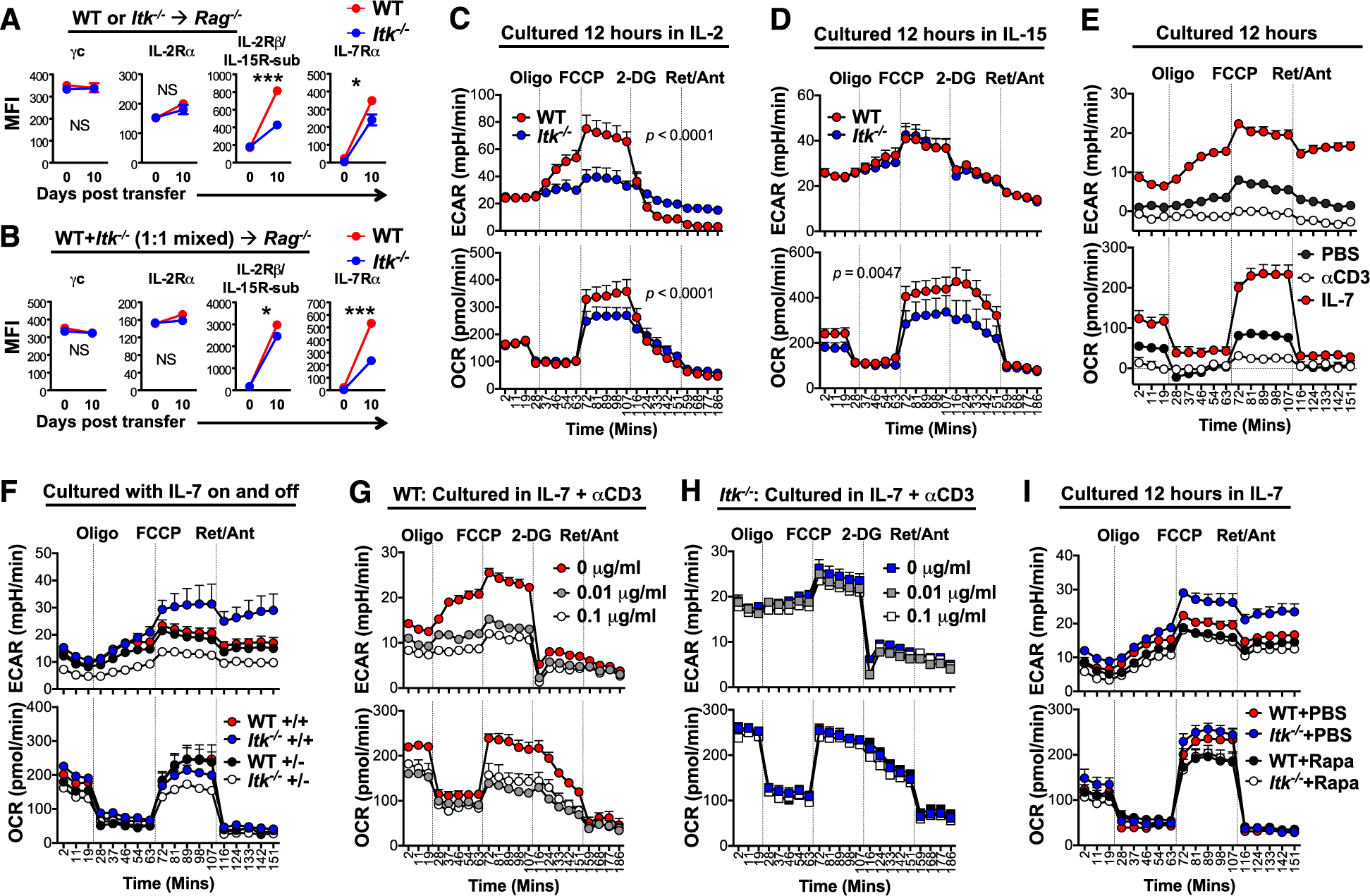
*Itk* deficient CD8^+^ T cells exhibit altered expression of γ chain cytokine receptors, metabolic potential, and are inert to TCR tuning. (A-B) Naive WT and *Itk*^−/−^ OTI-*Rag*^−/−^ CD8^+^ T cells are transferred singly (A) or in a 1:1 ratio mixed population (B) into *Rag*^−/−^ recipients day 0. Cells were analyzed for cytokine receptor expression on day 0 and day 10. MFI of the indicated cytokine receptors are shown. N = 3–7. (C-D) Naïve WT or *Itk*^−/−^ CD8^+^ T cells were cultured in Seahorse assay media with IL-2 (C) or IL-15 (D) for an overnight, and analyzed for ECAR and OCR. N = 4. (E) Naïve WT CD8^+^ T cells were cultured in Seahorse assay media with PBS, IL-7 or anti-CD3s (0.01 μg/ml) overnight, and analyzed for ECAR and OCR. N = 4–9. (F) Naïve WT or *Itk*^−/−^ CD8^+^ T cells were cultured in Seahorse assay media with continuous (+/+: media changed every 12 hours with IL-7 added each time) or intermittent (+/−: media changed every 12 hours between IL-7-supplemented and IL-7-free media) IL-7 for three days, and analyzed for ECAR and OCR. N = 4. (G-H) Naïve WT (F) or *Itk*^−/−^ (G) CD8^+^ T cells were cultured in Seahorse assay media with IL-7 in the presence of the indicated concentrations of anti-CD3s overnight, and analyzed for ECAR and OCR. N = 6–9. (I) Naïve WT or *Itk*^−/−^ CD8^+^ T cells were cultured in Seahorse assay media with IL-7 in the presence or absence of rapamycin (Rapa) overnight. N = 6. Data represent Mean ± SEM. *p < .05 by two-way ANOVA in the indicated sections.

**Figure S4.**
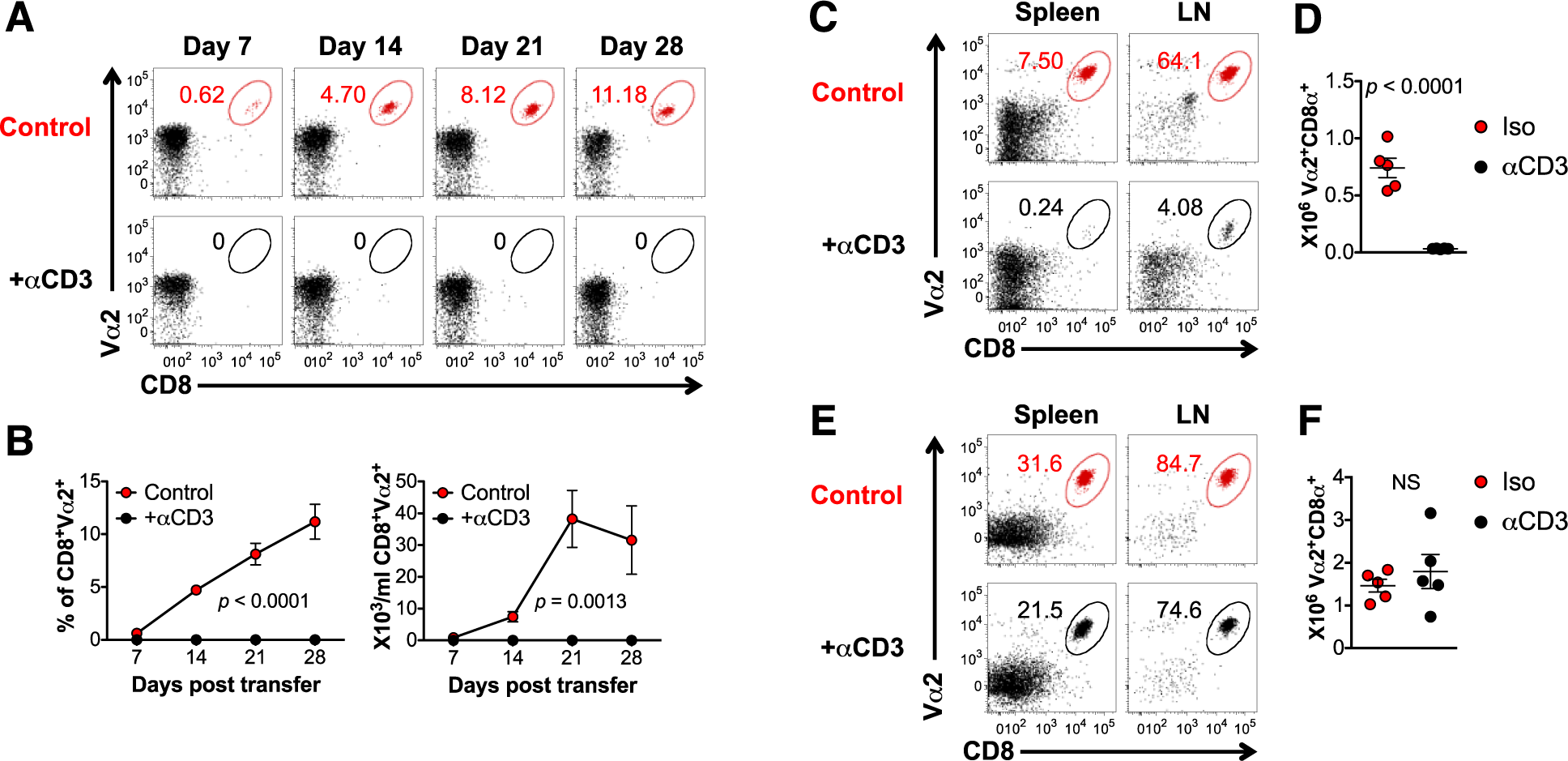
TCR activation in vivo reduces CD8^+^ T cell HP population in lymphopenic hosts. 120,000 naïve WT OTI-*Rag*^−/−^ CD8^+^ T cells are transferred into *Rag*^−/−^ recipients on day 0. (A-D) Recipients received intraperitoneal injection of 0.15 μg/mouse anti-CD3? (or isotype as control) on day 1 and day 3. Blood were sampled weekly as shown in (A&B), and spleen and lymph nodes were analyzed on day 30. N = 5. (A) Representative plots of WT CD8^+^ T cells in the blood following HP at the indicated time points. (B) Summary of percentage and number of HP CD8^+^ T cells over viable lymphocytes in the blood along the time shown. *p* values generated by two-way ANOVA. (C) Representative plots CD8^+^ T cells in the spleen and lymph node (LN) on day 30. (D) Total number of donor CD8^+^ T cells in the spleen and lymph node on day 30. *p* values generated by non-parametric Mann-Whitney test. (E-F) Recipients received intraperitoneal injection of 0.15 μg/mouse anti-CD3s (or isotype as control) on day 61 and day 63, and spleen and lymph nodes were analyzed on day 73. N = 5. (E) Representative plots of CD8^+^ T cells in the spleen and lymph node (LN) on day 73. (F) Total number of donor CD8^+^ T cells in the spleen and lymph node on day 73. *p* values generated by non-parametric Mann-Whitney test. Data presented as Mean ± SEM.

**Figure S5.**
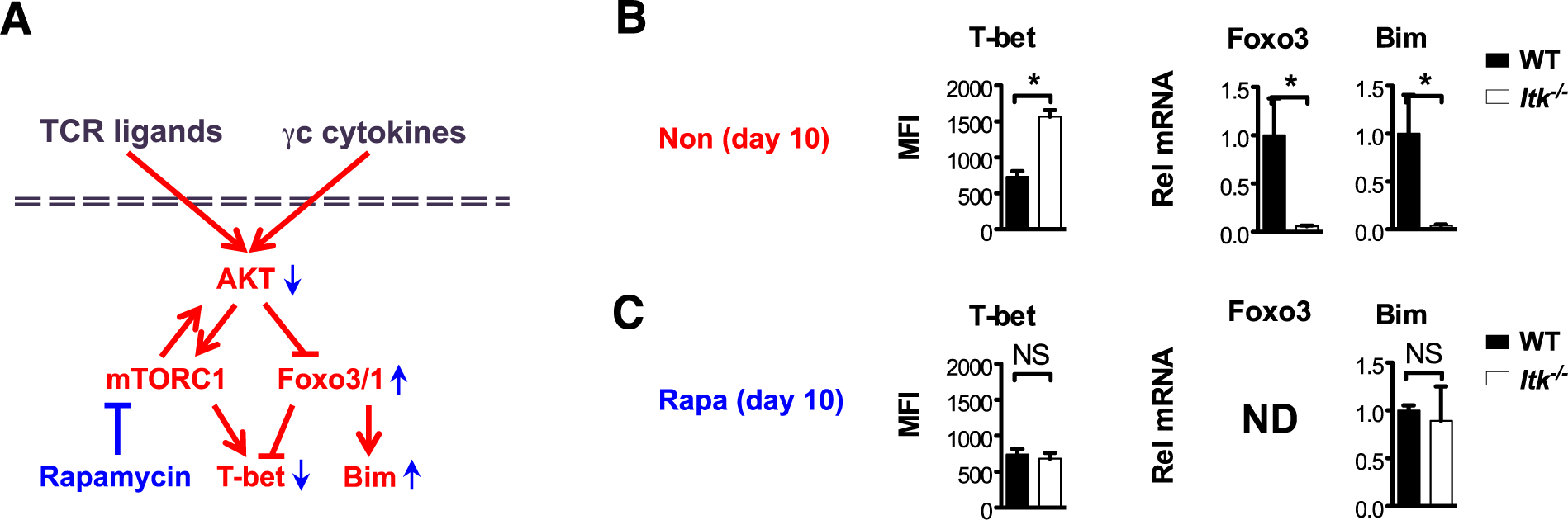
Inhibition of mTOR signaling reverts Bim and T-bet expression in *Itk*^−/−^ HP CD8^+^ T cells. (A) Model of TCR ligands and γc cytokines activation of mTOR signaling that suppresses Foxo3/Bim expression and enhances T-bet expression; inhibition of rapamycin can restore Foxo3-mediated Bim and T-bet expression (adapted from reviews (40, 41)). (B-C) 120,000 naïve WT or *Itk*^−/−^ OTI-*Rag*^−/−^ CD8^+^ T cells are transferred into *Rag*^−/−^ recipients on day 0, followed by PBS (B) or rapamycin (Rapa in C) treatment for 10 consecutive days, and cells pooled from spleen and lymph nodes were analyzed. (B) Expression level of T-bet (protein, MFI) is increased and levels of Foxo3a and Bim (mRNA) are reduced in *Itk*^−/−^ HP CD8^+^ T cells during early stage of lymphopenia-induced homeostatic proliferation. (C) Inhibition of mTOR by rapamycin reverted T-bet and Bim expression in *Itk*^−/−^ HP CD8^+^ T cells towards the WT levels (Foxo3 not detected (ND), likely due to low amount of HP CD8^+^ T cells expanded in the presence of rapamycin as shown in Figure 6H thus low amount of RNA in samples). Data represent Mean ± SEM. N = 3; \**p* < 0.05; NS = “No Significance” by Student’s *t* test.

**Figure S6.**
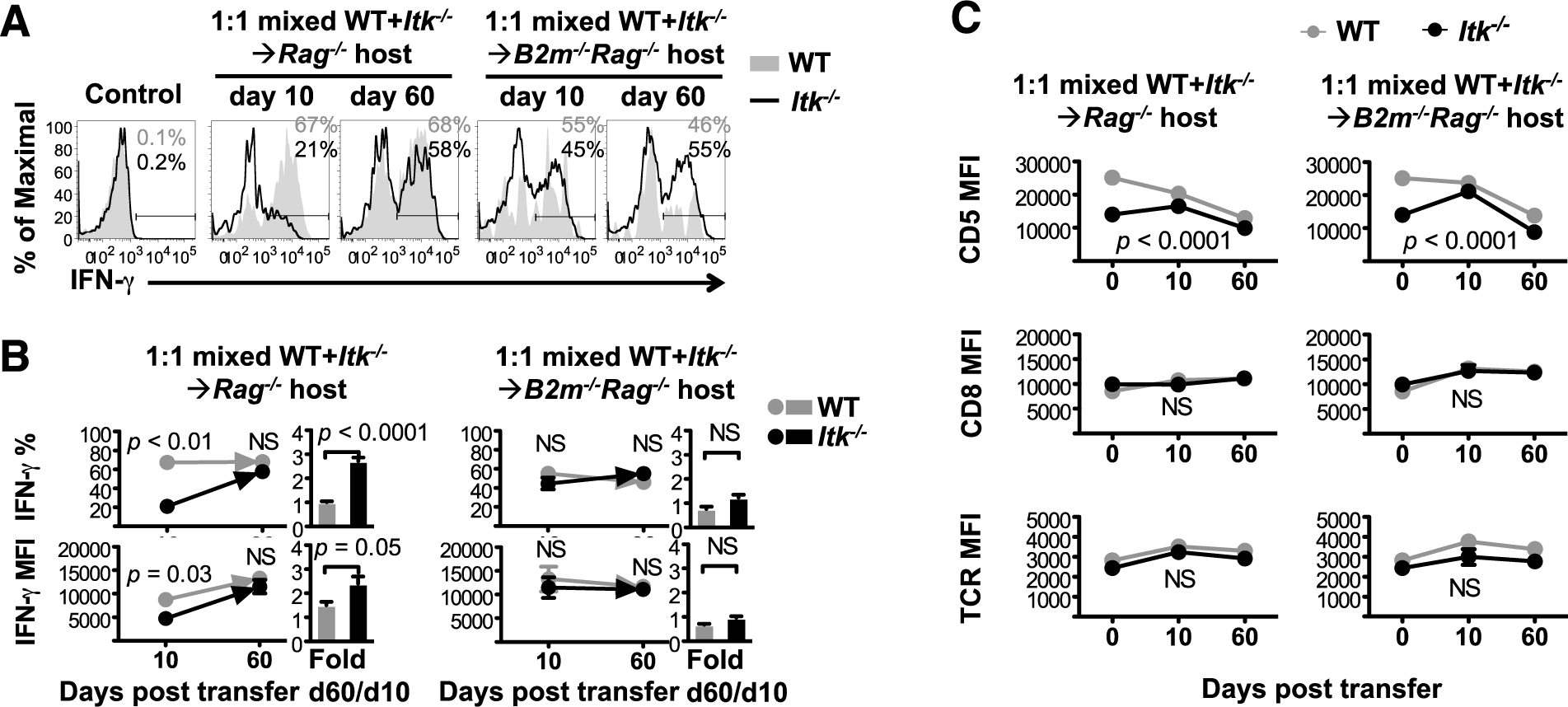
ITK regulates antigen sensitivity tuning during lymphopenia-induced homeostatic proliferation in CD8^+^ T cells. Naïve CD45.1^+^ WT and CD45.2^+^ *Itk*^−/−^ OTI-*Rag*^−/−^ CD8^+^ T cells (initial total number ≈ 0.1 × 10^6^ for each recipient) were expanded concurrently in *Rag*^−/−^ or *B2m*^−/−^*Rag*^−/−^ hosts for 10 and 60 days, followed by stimulation with OVA257–264 peptide in vitro. CD45.1^+^ WT and CD45.2^+^ *Itk*^−/−^ HP CD8^+^ T cells in the spleen of lymphopenic hosts were analyzed. (A) Representative plots of IFN-γ expression induced by OVA257–264 stimulation. Mock-stimulated (PBS/BFA) cells were used as background controls. Average percentage of IFN-γ producing cells shown on plot: WT in grey and *Itk*^−/−^ in black. (B) Percentage and MFI of IFN-γ producing HP cells. Fold changes were derived by dividing the levels on day 60 by those on day 10, in WT or *Itk*^−/−^ HP cells respectively. *p* values were generated by Student’s *t* test, comparing percentages at the same time point or fold changes connected. (C) Dynamic expression (MFI) of CD5, CD8a and TCR (Vα2) in WT or *Itk*^−/−^ CD8^+^ T cells during lymphopenia-induced HP. *p* values were generated by two-way ANOVA. N = 2 - 7. Data represent Mean ± SEM. NS = “No Significance”.

**Figure S7.**
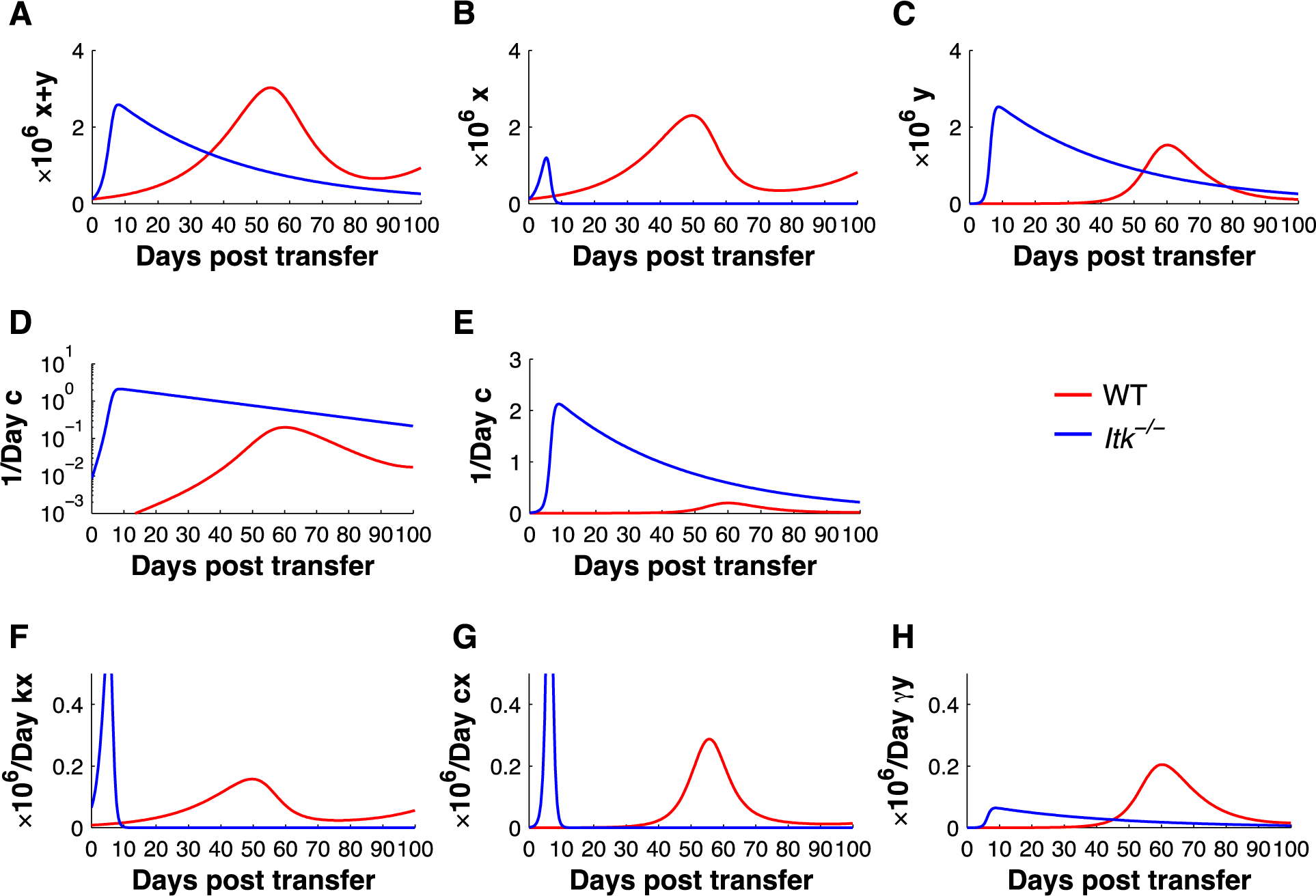
Simulated dynamics of. (A) Total HP CD8^+^ T cell population. (B) Population size of HP CD8^+^ T cell proliferating subset. (c) Population size of HP CD8^+^ T cell dying subset. (D) Conversion index in scientific notation. (E) Conversion index in linear form. (F) Proliferation rate. (G) Conversion rate from proliferating to dying subset. (H) Death rate.

**Table S1.**
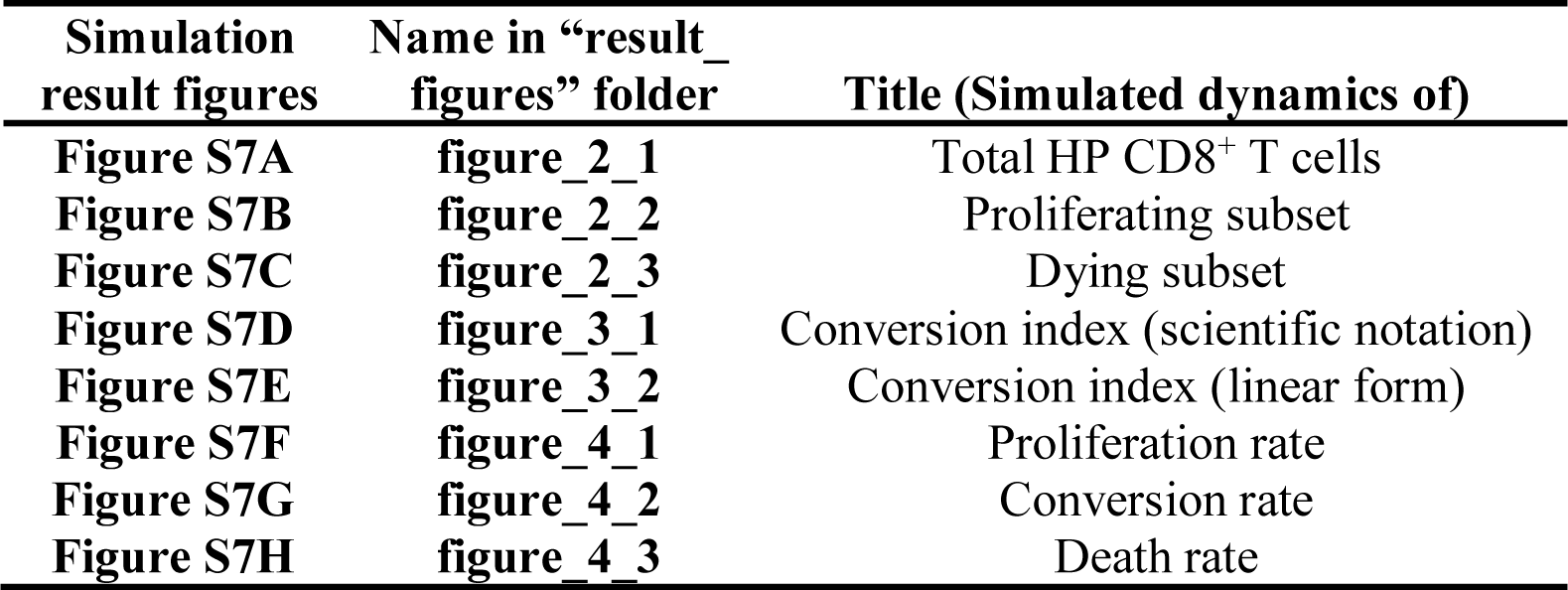
Annotation of values indicated by output figures in computational simulation results.

| **Simulation**<br>**result figures** | **Name in “result_**<br>**figures” folder** | **Title (Simulated dynamics of)** |
| --- | --- | --- |
| **Figure S7A** | **figure_2_1** | Total HP CD8<sup>+</sup> T cells |
| **Figure S7B** | **figure_2_2** | Proliferating subset |
| **Figure S7C** | **figure_2_3** | Dying subset |
| **Figure S7D** | **figure_3_1** | Conversion index (scientific notation) |
| **Figure S7E** | **figure_3_2** | Conversion index (linear form) |
| **Figure S7F** | **figure_4_1** | Proliferation rate |
| **Figure S7G** | **figure_4_2** | Conversion rate |
| **Figure S7H** | **figure_4_3** | Death rate |

